# Analysis of head and neck carcinoma progression reveals novel and relevant stage-specific changes associated with immortalisation and malignancy

**DOI:** 10.1101/365205

**Authors:** Ratna Veeramachaneni, Thomas Walker, Antoine De Weck, Timothée Revil, Dunarel Badescu, James O’Sullivan, Catherine Higgins, Louise Elliott, Triantafillos Liloglou, Janet M. Risk, Richard Shaw, Lynne Hampson, Ian Hampson, Simon Dearden, Robert Woodwards, Stephen Prime, Keith Hunter, Eric Kenneth Parkinson, Ioannis Ragoussis, Nalin Thakker

## Abstract

Head and neck squamous cell carcinoma (HNSCC) is a widely prevalent cancer globally with high mortality and morbidity. We report here changes in the genomic landscape in the development of HNSCC from potentially premalignant lesions (PPOLS) to malignancy and lymph node metastases. Frequent likely pathological mutations are restricted to a relatively small set of genes including *TP53, CDKN2A*, *FBXW7*, *FAT1*, *NOTCH1* and *KMT2D*; these arise early in tumour progression and are present in PPOLs with *NOTCH1* mutations restricted to cell lines from lesions that subsequently progressed to HNSCC. The most frequent genetic changes are of consistent somatic copy number alterations (SCNA). The earliest SCNAs involved deletions of *CSMD1* (8p23.2), *FHIT* (3p14.2) and *CDKN2A* (9p21.3) together with gains of chromosome 20. *CSMD1* deletions or promoter hypermethylation were present in all of the immortal PPOLs and occurred at high frequency in the immortal HNSCC cell lines (promoter hypermethylation ~63%, hemizygous deletions ~75%, homozygous deletions ~18%). Forced expression of *CSMD1* in the HNSCC cell line H103 showed significant suppression of proliferation (p=0.0053) and invasion *<u>in</u> <u>vitro</u>* (p=5.98X10^−5^) supporting a role for *CSMD1* inactivation in early head and neck carcinogenesis. In addition, knockdown of *CSMD1* in the *CSMD1*-expressing BICR16 cell line showed significant stimulation of invasion *<u>in vitro</u>* (p=1.82 x 10^−5^) but not cell proliferation (p=0.239). HNSCC with and without nodal metastases showed some clear differences including high copy number gains of *CCND1*, hsa-miR-548k and *TP63* in the metastases group. GISTIC peak SCNA regions showed significant enrichment (adj P<0.01) of genes in multiple KEGG cancer pathways at all stages with disruption of an increasing number of these involved in the progression to lymph node metastases. Sixty-seven genes from regions with statistically significant differences in SCNA/LOH frequency between immortal PPOL and HNSCC cell lines showed correlation with expression including 5 known cancer drivers.

**Lay Summary:** Cancers affecting the head and neck region are relatively common. A large percentage of these are of one particular type; these are generally detected late and are associated with poor prognosis. Early detection and treatment dramatically improve survival and reduces the damage associated with the cancer and its treatment. Cancers arise and progress because of changes in the genetic material of the cells. This study focused on identifying such changes in these cancers particularly in the early stages of development, which are not fully known. Identification of these changes is important in developing new treatments as well as markers of behaviour of cancers and also the early or ‘premalignant’ lesions. We used a well-characterised panel of cell lines generated from premalignant lesions as well as cancers, to identify mutations in genes, and an increase or decrease in number of copies of genes. We mapped new and previously identified changes in these cancers to specific stages in the development of these cancers and their spread. We additionally report here for the first time, alterations in *CSMD1* gene in early premalignant lesions; we further show that this is likely to result in increased ability of the cells to spread and possibly, multiply faster as well.

## Introduction

Globally, head and neck carcinomas account for over 550,000 new cases per annum with a mortality of approximately 275,000 cases per year (1). By far, the commonest site of cancer within this region is the oral cavity and the commonest type of tumour is squamous cell carcinoma (SCC), which accounts for over 90% of all malignant tumours at this site. HNSCC is associated with high mortality having an overall 5-year survival rate of less than 50%. Furthermore, both the disease and the multimodal treatments options involved are associated with high morbidity (2).

The molecular pathology of head and neck squamous carcinoma has been extensively studied previously and some of the common somatic genetic changes have been variably characterised (3-7). There have been some studies of the multistage evolution of these tumours (8) but this is less well characterised. A small number of tumours arise from pre-existing lesions (known as potentially malignant lesions or PPOLs) such as leukoplakia or erythroplakia, which display variable epithelial dysplasia (9). However, the vast majority are thought to arise *de novo* from macroscopically normal appearing mucosa or possibly undiagnosed PPOLs. Support for the latter comes from data showing that the transcriptional signatures of PPOLs are retained in unrelated samples of SCC both *<u>in vivo</u>* (10) and *<u>in vitro</u>* (11). Nevertheless, it is clear that tumours arise from within a wide field bearing the relevant genetic alterations and that there is a risk of synchronous or metachronous tumours (12), (13). A fuller understanding of the events in evolution of these cancers may permit the development of biomarkers or effective therapeutic interventions possibly targeting not just tumours but also the early changes in the field. The first multi-step model proposed for carcinogenesis in HNSCC (14) suggested typical alterations associated with progression from normal mucosa to invasive carcinoma, with dysplasia reflecting an earlier stage of cancer progression.

We, and others, have previously shown that both SCCs and PPOLs yield either mortal and immortal cells *<u>in vitro</u>* (15), (16) and, sometimes, mixtures of the two (15), (16), (17). The status of these mortal cells is unclear. Unlike the immortal cells, they lack inactivation of *TP53* and *CDKNA,* but our limited previous investigations show that they are genetically stable. Nevertheless, they often have extended replicative lifespans (15, 16), possess neoplastic phenotypes, such as resistance to suspension-induced terminal differentiation (15) and have expression signatures which are distinct from both immortal cells and normal cells (11). Furthermore, these characteristics are present in mortal cells from both PPOLs and SCCs (11) suggesting the presence of distinct pathways for the development of mortal and immortal SCC. Our preliminary work established that the mortal PPOL were cytogenetically diploid and had low levels of LOH (15) but the immortal PPOL and SCCs have never been subjected to extensive genomic analysis. In addition, whilst extensive genetic analysis of HNSCC has been carried out in recent years (Stransky 2011; Agrawal et al 2011; Pickering et al 2013, The Cancer Genome Network 2015), including the identification of key driver mutations, the stages in the cancer progression, at which they occur and the resulting phenotypes are still unknown. There are numerous previous studies of PPOLs limited to determining the frequency of alterations in specific gene or specific genetic regions. Exceptions to this are the study by Bhattacharya and colleagues (8) and Wood and colleagues (18), which reported a comprehensive analysis of copy number variation in primary PPOLs and HNSCC although the later study, the PPOL analyses was confined to metachronous lesions. Here we extend their findings by a combination of exome/targeted sequencing and SNP/CGH array analyses, using our unique panel of mortal cultures and immortal cell lines derived from both PPOLS and HNSCC, to show that mostly these genetic alterations are mostly associated with cellular immortalisation and increase with the stage of tumour progression in this class of SCC keratinocyte.

## Results

### Mutation analyses

Several recent genomic sequencing studies have fully characterised mutations in HNSCC (3),(4), (5), (6). In order to map these previously identified mutations to progression of HNSCC, a small previously well-characterised panel of samples consisting of 3 PPOL mortal cultures, 7 PPOL cell lines (from progressing and non-progressing lesions) 1 mortal culture derived from HNSCC and 11 HNSCC cell lines (16, 19, 20) were selected for exome-sequencing or targeted sequencing of the top 40 genes identified as altered in these cancers (3) using HaloPlex Target Enrichment System (Agilent, Santa Clara, CA, USA). The sample details are given in Fig. 1 and in S1 Tables.

**Fig 1.**
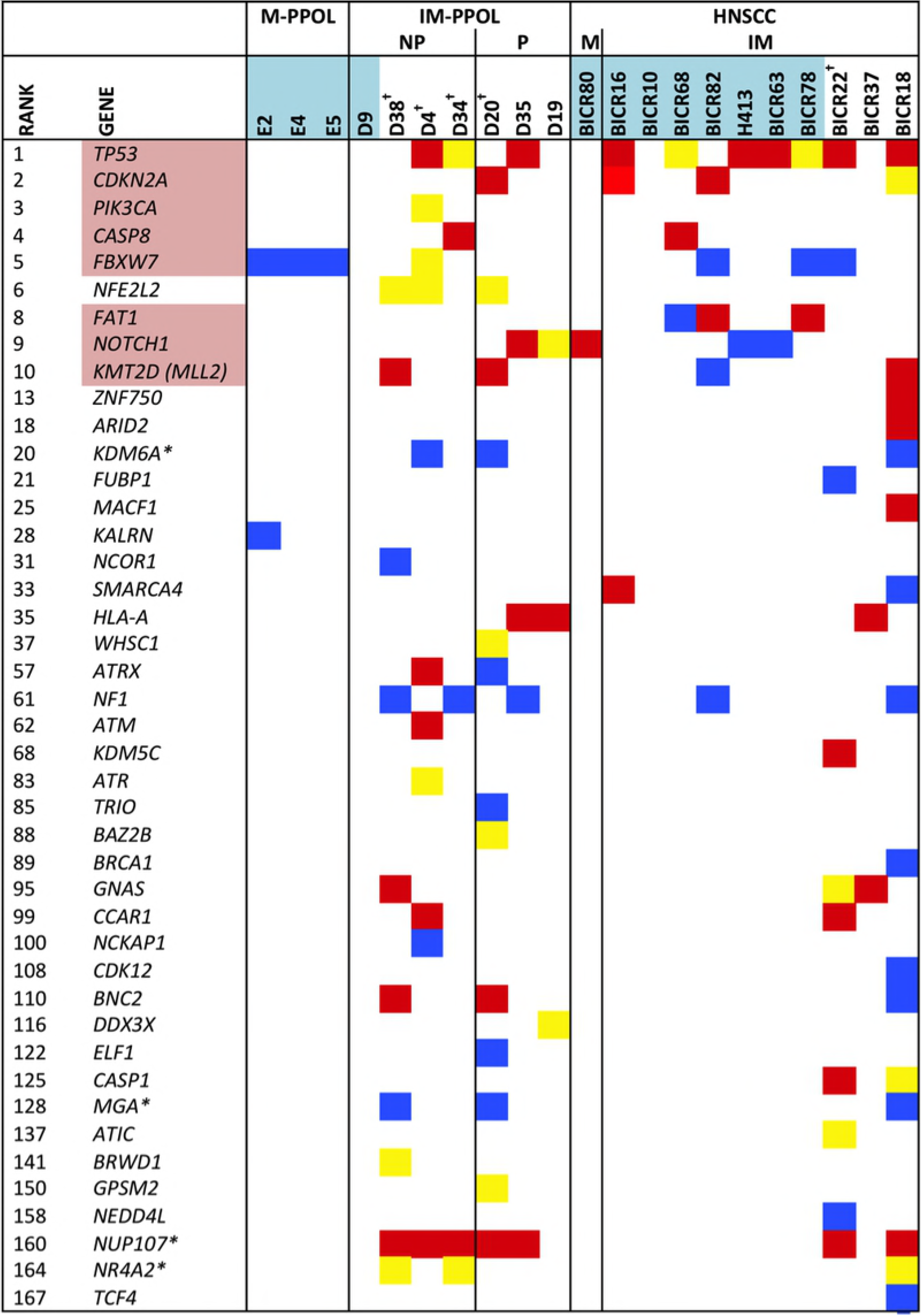
Mutations of IntOgen (version 2014.12; http://www.intogen.org) - predicted head and neck cancer driver genes in mortal PPOL (M-PPOL) cultures, immortal PPOL (IM-PPOL) progressive and non –progressive (P and NP respectively) cell lines and mortal HNSCC cultures (M) and immortal (IM) cell line panels. The green shaded sample number indicate samples that were subject to HaloPlex target enrichment and sequence analyses of specific genes The remainder were subject to exome sequencing. The gene names shaded indicate genes in the HaloPlex panel that are also HNSCC cancer drivers as indicated by IntOgen. Samples with at least one mutation predicted to be high impact (nonsense mutations, indels, frameshift, splice site) are shown in red, samples with at least one missense mutation predicted to be deleterious by both Polyphen and SIFT are shown in yellow, and samples with at least one missense mutation predicted to be deleterious by Polyphen or SIFT (but not both) are shown in blue. Any variants previously reported as constitutional SNP was excluded regardless of the minor allele frequency unless it had been demonstrated to be pathogenic previously. The ranking is from IntOgen and indicates ranking by frequency of mutations in HNSCC. *Indicates a gene with the same variant in each sample with a variant. ^†^ Indicates samples with low coverage (<80% x20) in exome sequencing. Only genes showing significant mutations by criteria used are shown here.

For exome sequencing, approximately 6 gigabases of sequence mapped to the human genome with an average of 65.7% (Range 33.8% to 86.1%) of the targeted exome covered at twenty-fold or higher (S2 Fig). The lower coverage was sample specific and these samples are indicated in Fig. 1. For HaloPlex sequencing, approximately 800 megabases of sequence mapped to the human genome with an average of 94.8% of the targeted exons covered at twenty-fold or higher (S2 Table).

For calling pathological mutations, we used the strategy described in detail in the legend for Fig. 1; this was stringent and therefore, it is possible that some genuine pathological mutations may have been excluded. However, the full dataset with GEMINI framework annotation (21) and raw data is available through Dryad Digitial Repository (URL: https://datadryad.org/review?doi=doi:10.5061/dryad.314k5k5 Provisional doi:10.5061/dryad.314k5k5)

Given the small numbers of samples examined in our study, we targeted our analyses to 167 cancer drivers in head and neck cancers as identified by IntOGen (Release 2014.12) The HaloPlex sequencing panel included 8 of the 10 most frequently mutated HNSCC driver genes. Limiting analyses to known cancer drivers also effectively excluded possible false positives that can arise due to DNA replication timing and low transcriptional activity (22). Thus, our significant driver mutations (Fig. 1) largely mirror those identified by Lawrence and colleagues, 2013 (22) following correction for these factors. Significant mutations are shown schematically in Fig. 1 and detailed in S 2C-2D Tables.

Mutations were rare in mortal cultures (Fig. 1). A single missense variant of *FBXW7* predicted to be deleterious with low confidence was observed in all 3 PPOL cultures (in addition to several immortal HNSCCs) and one high impact *NOTCH1* mutation was observed in HNSCC culture BICR80.

As with previous studies (4), (3), (5), (6), the mutation analyses revealed a small set of genes (*TP53, CDKN2A, FBXW7, FAT1, NOTCH1* and *KMT2D*) as the most common targets for likely deleterious sequence mutations in immortal PPOL and HNSCC cell lines (Fig. 1). Mutation of *TP53* and *CDKN2A* as an early event in head and neck carcinogenesis is well established (23), (24), (16), (25). In the present study, however, we demonstrate for the first time that mutations in the other commonly mutated genes in HNSCC also occur early and are not only present in PPOLs derived from progressing lesions (D19, D20, D35) but also in non-progressing lesions (D4, D34, D38).

As SCNAs may be an alternative mechanism for gain or loss of function of a gene, we examined the frequency of SCNAs of genes predicted to be cancer drivers by IntOgen but not showing sequence variation, in our panel. Many cancer drivers showed a high frequency of SCNAs (S3 Tables). These included well-characterised tumour suppressors and oncogenes implicated in HNSCC such as *CDKN2A*, *CCND1*, *EGFR, PIK3CB* but also other cancer drivers that are less well characterised in HNSCC (e.g., *CTTN*, *NDRG1, MLL3, ROBO2*). In addition, there were clear differences between PPOLs and HNSCCs as well as between cell lines from HNSCC with and without lymph node metastases. Although it can be argued that some of these SCNAs may be bystander changes in chromosomal gains or losses targeting other genes, consistent high frequency changes are likely to be important as indicated by inclusion of already well-characterised head and neck cancer drivers such as *MYC*, *PIK3CA* and *CDKN2A*.

### Early changes in evolution of HNSCC

Somatic copy number alterations in 7 PPOL cell lines, 11 mortal cell cultures derived from PPOL, were identified using Illumina HumanHap550 Genotyping Beadchip and Infinium Assay II. The full dataset including raw data and Nexus Copy Number v5.1 (BioDiscovery, Inc., CA, USA) data are available at Dryad Digital Repository (URL: https://datadryad.org/review?doi=doi:10.5061/dryad.314k5k5 Provisional doi:10.5061/dryad.314k5k5)

#### Mortal cultures are genetically stable

Mortal cultures derived from PPOL were genetically stable and showed very few copy number changes, data that are consistent with their diploid chromosome complement and previous limited loss of hetrozygoisty analysis (15) (Fig 1 and S4 Fig.). There were no statistically significant differences (P>0.05) in SCNA between mortal PPOLS and matched fibroblasts.

One mortal cell culture from PMOL (D17) that has an extended lifespan and does not express CDKN2A but regulates telomerase normally and has a functional *TP53* gene (11, 16, 19). This cell culture showed deletion of one chromosome arm 9p with duplication of the homologous chromosome 9, 3 copies each of chromosome 2 and 7 and uniparental trisomy of chromosome 5 (data not shown). It is possible that the changes involving 9p reflect loss of a normal *CDKN2A* allele and duplication of an allele with hypomorphic mutation.

#### Immortal PPOL cell lines show progressive changes principally involving chromosome 3, 8, 9 and 20

Immortal cultures derived from PPOLs showed consistent SCNAs, thereby defining the earliest changes in the development of HNSCC (Fig. 2a). Overall, there were statistically significant (P<0.05) losses on chromosome 3p, 8p and 9p coupled with gains of chromosome 20 compared to normal fibroblasts (Fig. 2b). However, there was clear heterogeneity in the nature of the changes observed in the 7 immortal PPOL samples (Fig. 2c) with some specimens showing more frequent changes than others. Meaningful testing for significant differences between subsets of PPOLS was not possible due to the small sample size and thus, hierarchical clustering was used to group samples by SCNAs (Fig. 2c). This was remarkably similar to the clustering previously observed for these cell lines by gene expression profiles (11). The samples clustered into two main groups that suggested a correlation of genomic changes with grade of dysplasia; the two cell lines derived from PPOLs with mild dysplasia clustered separately from those derived from PPOLs with higher-grade dysplasia. Furthermore, in the latter group, the three cell lines (D19, D20, D35) from PPOLs that progressed to carcinomas (progressive PPOLs or P-PPOL), clustered in a discrete subgroup distinct from cell lines (D34, D4) derived from non-progressive lesions (N-PPOL). Distinct gene expression signatures for normal tissues, PPOLs and HNSCCs have been reported previously (10). In the present study, we provide evidence that these signatures may at least in part reflect the underlying somatic copy number changes.

**Fig 2.**
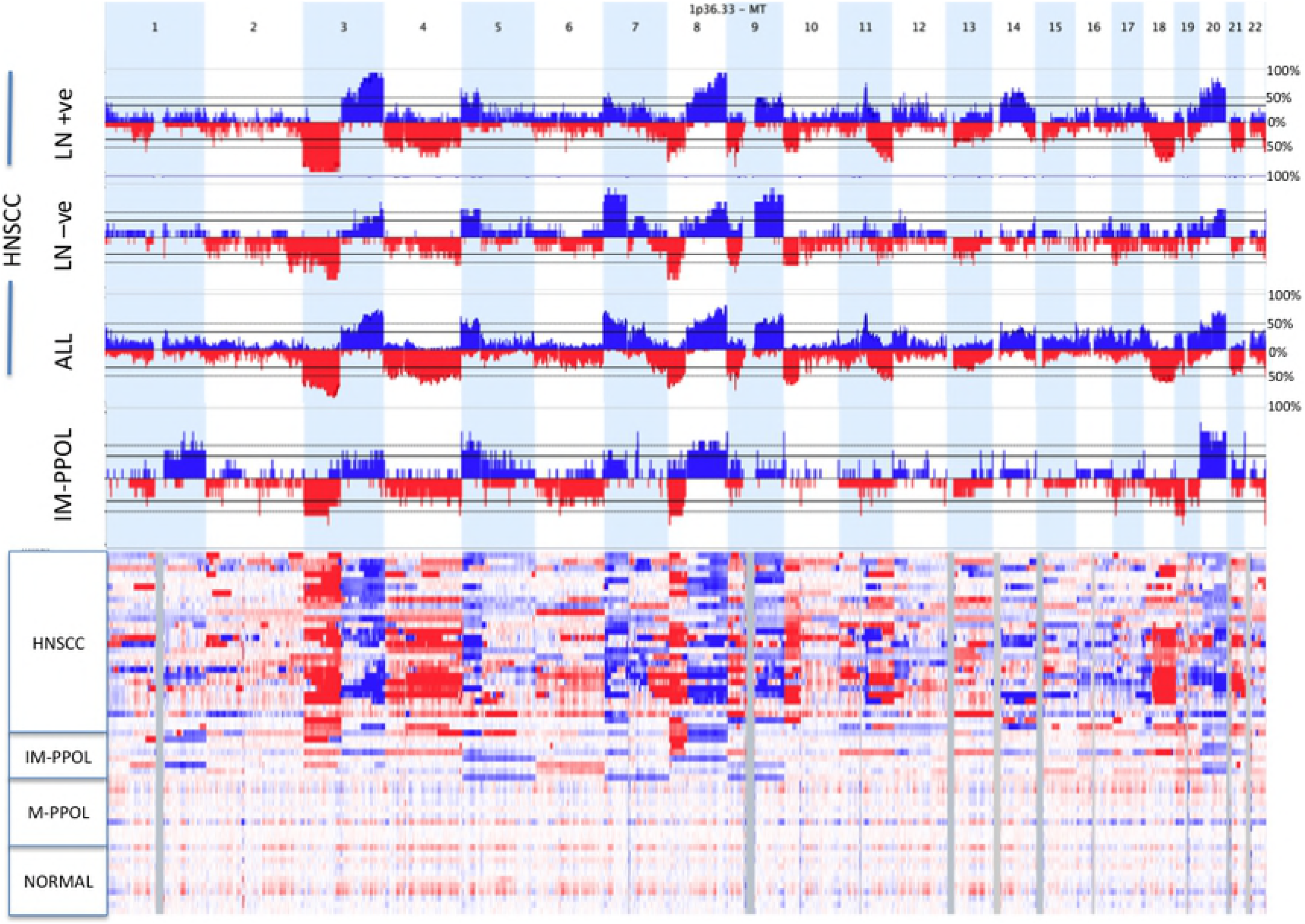
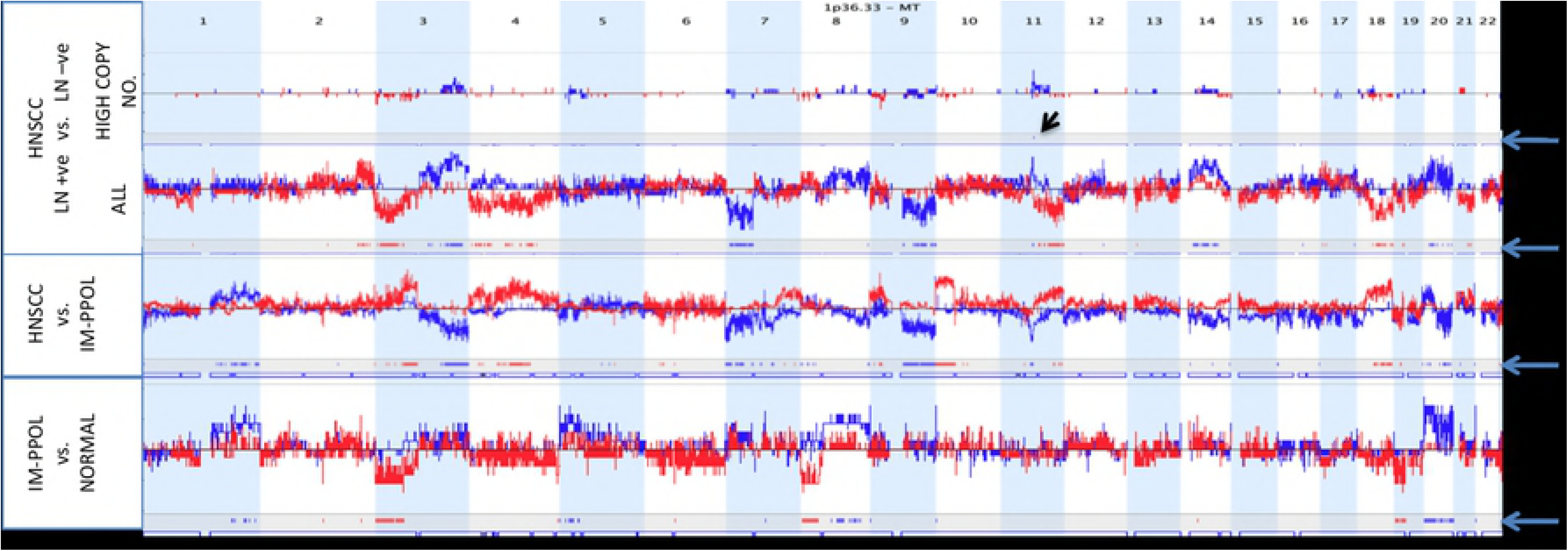
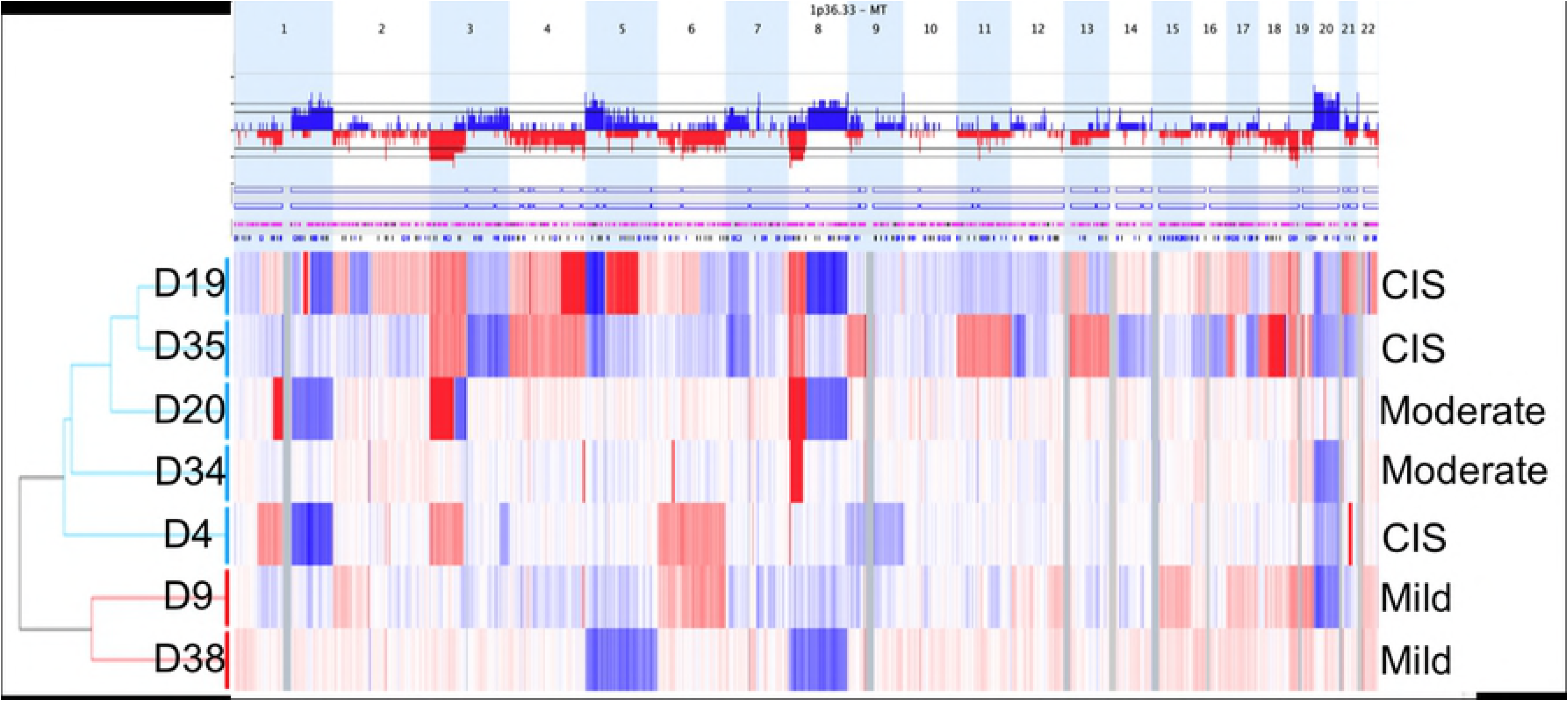
Somatic copy number changes in HNSCC. **A**. Density plots and karyograms showing copy number gains (blue) and losses (red) in normal fibroblasts, mortal PPOLs (M-PPOL), immortal PPOLs (IM-PPOL) and HNSCC cell lines with and without lymph node metastases (LN+ve and LN-ve respectively). **B.** Subtraction karyograms showing differences in copy number gains (blue) and losses (red) between (i) normal fibroblasts, and mortal PPOLs (M-PPOL), (ii) immortal PPOLs (IM-PPOL) and all HNSCC cell lines and (iii) HNSCC cell lines with and without lymph node metastases (LN+ve and LN-ve respectively). Both high and all copy number changes are shown for LN+ve and LN-ve HNSCC cell lines. Blue arrows on the right indicate lanes with regions of difference showing statistical significance (p<0.05). **C**. Density plots and karyograms showing copy number gains (blue) and losses (red) in individual immortal PPOL cell lines showing hierarchical clustering and grade of dysplasia.

Cell lines from progressive lesions (D19, D20, D35) were characterised by consistent arm-level losses of chromosome arms 3p and 8p with homozygous deletions of *FHIT* (3p14.1) and *CSMD1* (8p23.2) (S5 Fig). In addition, two of the three P-PPOLs showed further arm-level SCNAs on several other chromosomes (+3q, +5p, +7p +8q, −13p, −13q, −18p −18q, +20). The pattern of SCNAs in cell lines derived from lesions that had not progressed to date (NP-PPOLs, D34, D4, D9 and D38) reflected their earlier stage of evolution (Fig. 2c). These cells harboured focal SCNAs involving the *CSMD1* (3 of 4 cell lines) and *FHIT* (2 of 4 cell lines) on chromosome 8p23.2 and 3p14.1 respectively, and showed arm-level gains of chromosome 20 (3 of 4 cell lines). Additionally, other chromosomal arms (+3q, +5p, +7p +8q, −13p, −13q, −18p −18q) also showed variable and largely focal SCNA.

#### GISTIC analyses identifies key genes deleted early in progression of HNSCC

Significant peaks of copy number gains and losses were identified using GISTIC (26). The genes in the peak regions at different levels of stringency (Q=0.05-0.25) are shown in S6 Table. These included the homozygously deleted genes described above (*FHIT*, *CSMD1*, *CDKN2A, CDKN2B*). In addition, the peaks included homozygous deletions of *FAT1*, thereby supporting loss-of-function mutations in HNSCC and identifying inactivation of *FAT1* as an early change in HNSCC development. Homozygous deletions of other genes (*NCKAP5, SORBS2, FAM190A)* not previously implicated in HNSCC were also identified in peak regions. Some genes such as *PTPRD, LRP1B* and *LINGO2*, which are target for homozygous deletions in HNSCC cell lines, showed frequent hemizygous deletions in the PPOL cell lines suggesting further selection in progression to HNSCC. Deletion of *NCKAP5* was explored a little further (S7) as rare SCNAs involving *NCKAP5* have been reported recently in HNSCC (3) (in supplementary data) and in prostate cancer (27). Potentially deleterious mutations have been reported in COSMIC (S7 Table). *NCKAP5* homozygous deletions were present in both immortal PPOL and HNSCC cell lines and the homozygous deletions eliminated one or more exons in 4 out of the 5 cell lines (S7 Fig) with the remaining cell line sustaining two intronic deletions. However, deletions of *NCKAP5* are not common in primary HNSCC and other tumour types (Tumorscape Release 1.6, our tumour panel) and furthermore expression was not reduced significantly in HNSCC and other tumour types (S7 Fig). Thus, *NCKAP5* is unlikely to be frequent target for inactivation in HNSCC but the related pathways may be more significant. NCKAP5 has been shown to interact with NCK1 (STRING Release 9.1), an adaptor protein important in ligand-induced activation of receptor tyrosine kinases (28), and also APC (BioGRID Release 3.3). Our IntOGene (Release 2014.12) analyses showed a low frequency of pan-cancer gain-of-function mutations in *NCK1*. In this study, *NCK1* was within an extended GISTIC region and copy number gains were seen in 60% of LN+ve cell lines with relatively low frequency (~14%) gains in PPOL and LN-ve cell lines.

#### CSMD1 shows SCNAs early in development of HNSCCs and functional analyses suggest a tumour suppressor role

Inactivation of *TP53* and *CDKN2A* are almost universal in both P-PPOL and NP-PPOL immortal cell lines (16) and in HNSCC in vivo unless integrated oncogenic HPV is present (7). Our data indicate that specific focal deletions of *FHIT* and *CSMD1* are also common early changes. *FHIT* is already well characterised and appears to have a tumour suppressor function (29, 30) but relatively little is known about *CSMD1*. Thus, we explored whether *CSMD1* functions as a tumour suppressor in HNSCC. *CSMD1* is deleted in many tumour types (31) and also shows rare somatic mutations (32). We observed both homozygous and hemizygous deletions of *CSMD1* (5/28 and 21/28 respectively) in HNSCC cell lines (S8 Fig). Analysis of expression in ProteomicsDB (33) suggests that the highest expression in the body is in the oral epithelium although generally the high expression is reported in brain and testis (Gene, NCBI).

Only one immortal PPOL (D4) sustained a nonsense mutation (G1579X) in *CSMD1* and no likely pathogenic mutations were observed in HNSCC. *CSMD1* promoter methylation analyses by pyrosequencing revealed hypermethylation in 9 of 12 (75%) HNSCC cell lines with matching normal samples (and several other HNSCC cell lines without matching normal tissues), 3 of 7 (~43%) PPOL cell lines, and 15 of 24 (~63%) primary HNSCCs (Figure 2). The highest level of promoter hypermethylation was seen in immortal HNSCC cell lines with either hemizygous deletions (BICR56, BICR22, BICR82, BICR10 and T4) or absence of deletions (BICR63, BICR68 and H314), and in the only immortal PPOL cell line (D9) that lacked *CSMD1* deletions (Fig. 3, and S5 and 8 Fig). Additionally, the frequency of promoter methylation in primary tumours (~63%) was not too dissimilar to the frequency of hemizygous deletions (~75%) in our HNSCC cell lines. These findings suggest that frequent inactivation of *CSMD1* in HNSCC occurs by deletion and/or promoter hypermethylation.

**Fig 3.**
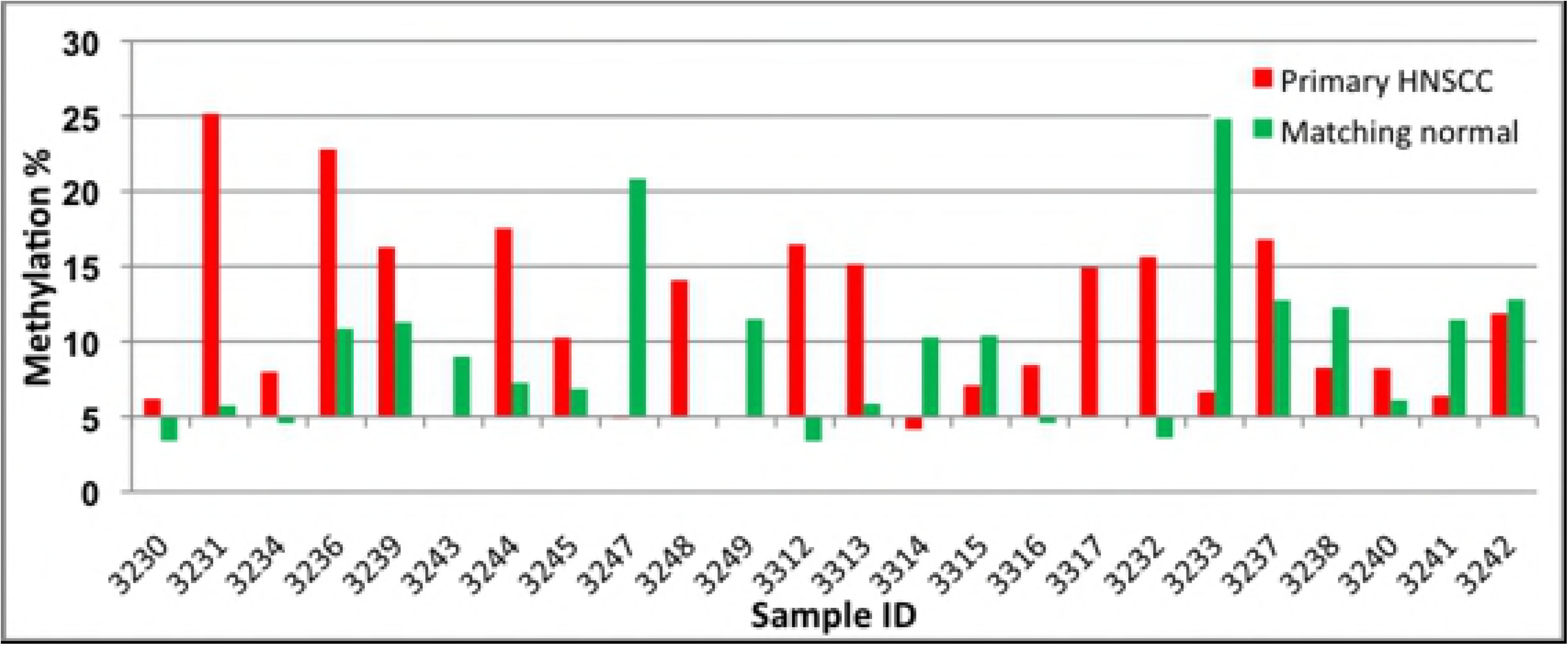
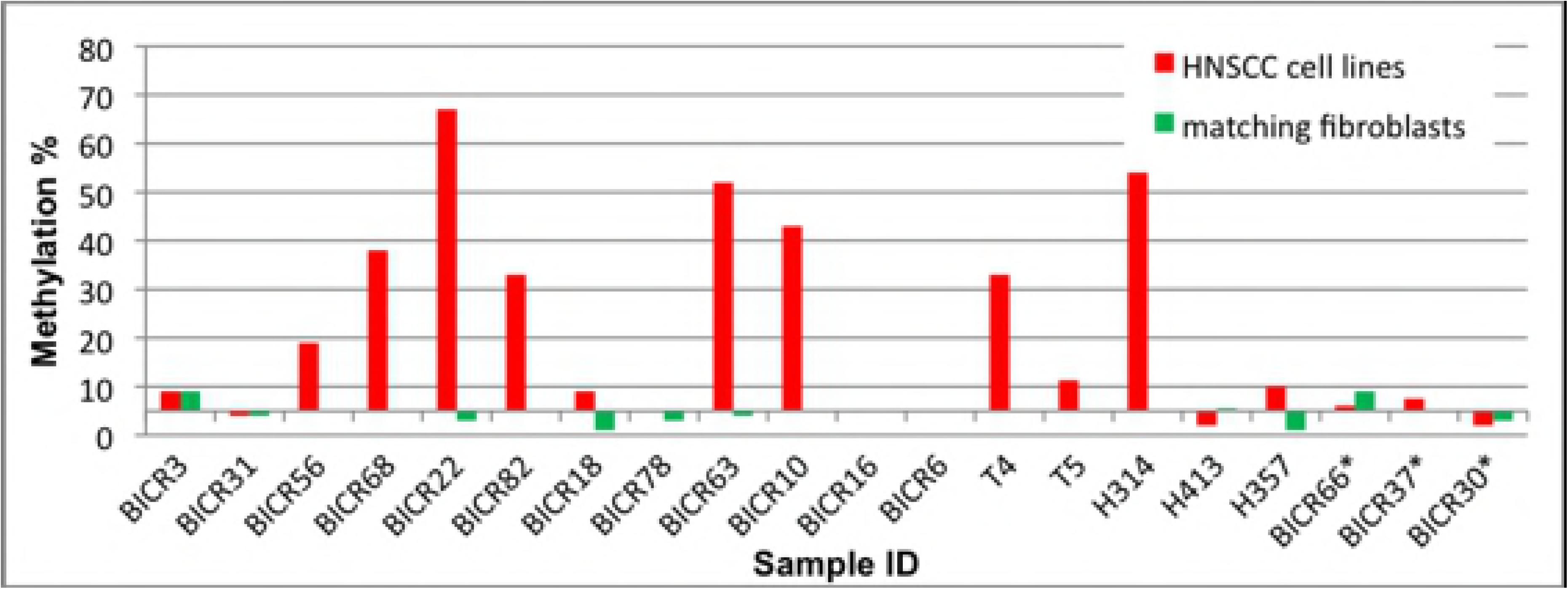
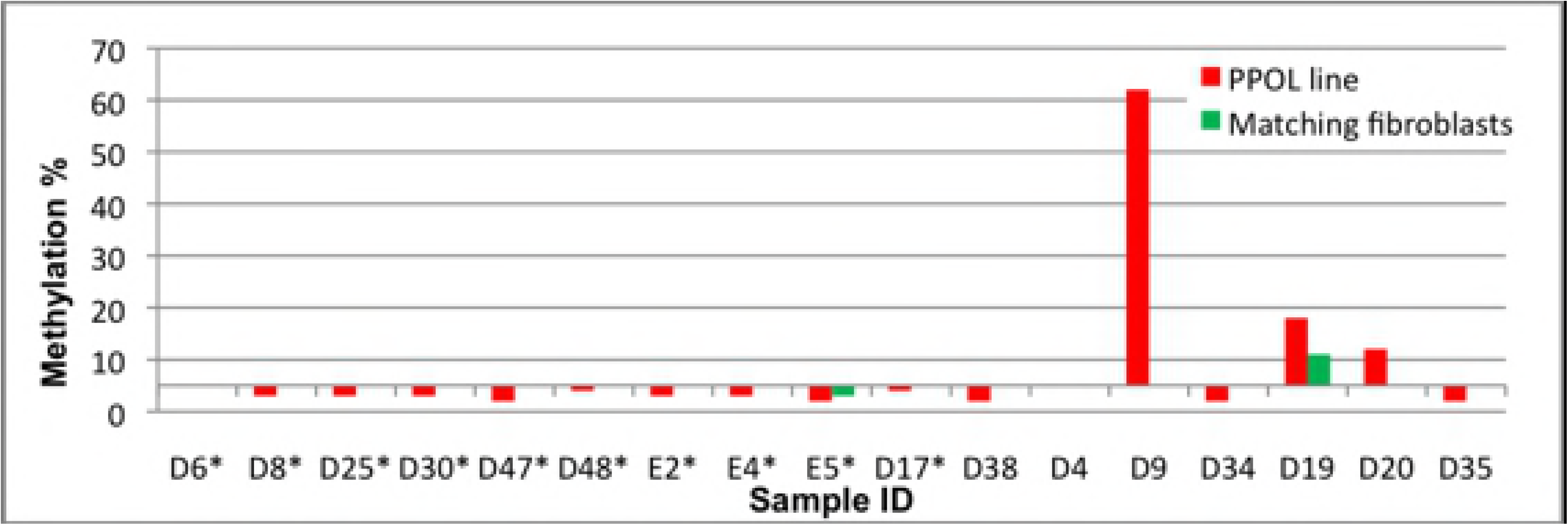
Clustered column chart representing pyrosequencing analyses of the *CSMD1* promoter region. Sample ID is shown on the X-axis and mean methylation percentage is represented on the Y-axis. For each sample, a mean methylation percentage greater than 5% was considered as significant promoter methylation. **A.** Primary HNSCCs and matching normal tissues. Significant promoter methylation was observed in 15 of 24 (~63%) primary HNSCC. **B.** Mortal and immortal HNSCC cell lines. Significant promoter methylation was observed in12 of 17 (~70%) immortal HNSCC cell lines but not in any of the mortal HNSCC cell lines (indicated by *). The highest levels of promoter methylation were in cell lines with hemizygous deletions (BICR56, BICR22, BICR82, BICR10 and T4) or no deletions (BICR63, BICR68 and H314). **C.** PPOL cell lines/cultures. Significant promoter methylation was observed 3 of 7 (~43%) immortal PPOL lines but not in any of the mortal PPOL cultures (indicated by *). The highest level of promoter methylation was in a single immortal cell line (D9) that had no deletions at the *CSMD1* locus.

Stable transfection of full-length *CSMD1* cDNA into the H103 cell line, which lacks endogenous expression of *CSMD1* expression, resulted in a significant inhibition of proliferation (p=0.0053) and invasion (p=5.98X10^−5^ - Matrigel *<u>in vitro</u>* assay). Results from a representative clone are shown in Fig. 4 (with data from all clones shown in S9 Fig). *CSMD1* expression was also silenced by stable transfection with an shRNA vector in the BICR16 cell line, which has a very low but detectable level of endogenous *CSMD1* expression. This cell line has complete hemizygous deletion of the *CSMD1* together with several small homozygous deletions of which one involves an exon (S8 Fig) although the functional state of the protein is unknown. The stable *CSMD1*-silenced clones showed a more variable effect on both proliferation and invasion; clones displayed a significant increase in invasion compared to parent cells (p = 1.82 x 10^−5^) but loss of *CSMD1* did not have a significant effect on the rate of proliferation (p = 0.239). Data from representative clones are shown in Fig. 4 (with data from all clones show in S9 Fig). It is possible that the inconsistent proliferation results with gene silencing are due to the fact that these cell lines have acquired the necessary cancer traits with some CSMD1 expression and that these traits are not significantly impacted by additional knockdown of residual and possibly hypofunctional CSMD1. Overall, however, these data support a role for *CSMD1* as a tumour suppressor gene inactivated in the very early stages of HNSCC development.

**Fig 4.**
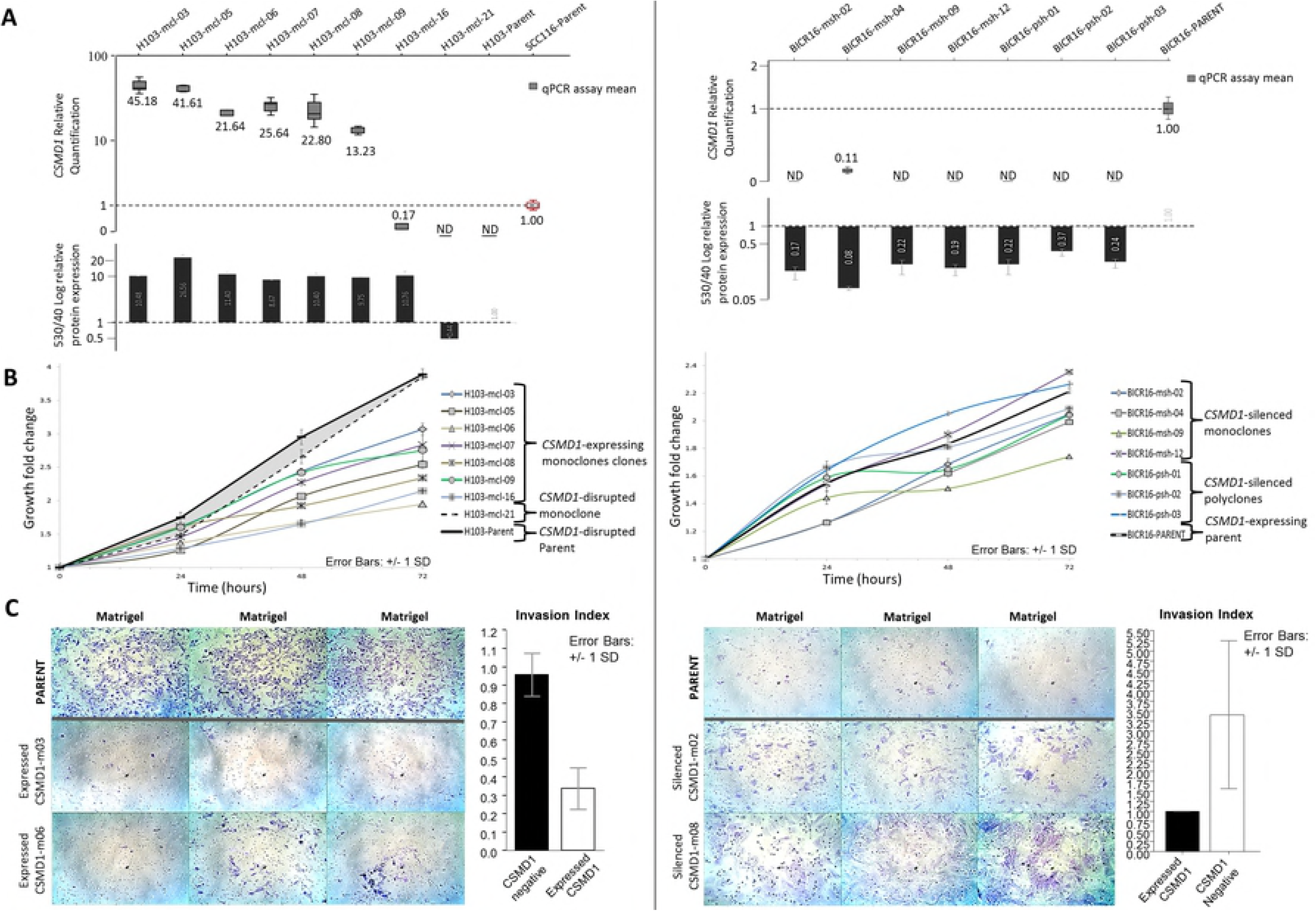
Phenotypic effects of *CSMD1* expression modulation in HNSCC. Left panel: forced *CSMD1* expression in the *CSMD1* non-expressing H103 cell line. Right panel: silencing of *CSMD1* in the *CSMD1-*expressing BICR16 cell line. A. *CSMD1* mRNA transcript quantification by RT-qPCR and protein quantification by flow cytometry for generated clones, presented as fold change normalised to the reference cell line. Left: H103 *CSMD1*-expressing clones and H103 *CSMD1*-negative parent cells and *CSMD1*-disrupted clone H103-mcl-21, normalised to SCC116 *CSMD1* expressing cells (red outline). Right: BICR16 *CSMD1*-silenced clones normalised to *CSMD1*-expressing BICR16 parent cell line. Boxplots represent RQ normalised to reference cells. Each box plot is the relative quantification (RQ) of two plates each of triplicate target and reference gene CT values plus and minus log-transformed standard deviations and so incorporates intra-plate variance. Standard boxes depict the first-third quartiles, whiskers depict ±1.5IQR. Median values are provided. Bar charts represent CSMD1 protein fold change normalised to reference cells. Error bars are ± standard deviation. B. Effects of modulation of *CSMD1* expression on cell proliferation. Left: *CSMD1*-expressing H103 monoclones compared to *CSMD1*-negative H103 parent and control cells. *CSMD1* expression resulted in a reduced growth rate compared to *CSMD1*-negative parent (black line) and H103-mcl-21 cells (dotted black line) (shaded area) (p = 0.0053). Right: Cell proliferation of *CSMD1-*silenced BICR16 clones compared to *CSMD1*-expressing BICR16 parent cells. *CSMD1* silencing did not result in a significant change in growth rate compared to *CSMD1*-expressing parent cells (black line) (p = 0.239). This observation was further confirmed using an additional *CSMD1-*silenced OSCC cell line with 11 silenced monoclones and 1 silenced polyclone (data not shown). Plots represent triplicate points from duplicate 96AQ assays for 96 hours with growth rates normalised to achieve relative fold change values for 0-72hrs. Error bars are ± 1 SD. Growth rate differences across the BICR16 clones and parent cells illustrate the proliferation variance inherent across the clone pool, rather than differences being specifically due to loss of expressed CSMD1 underpinning a clonal growth variation effect. C. Effects of modulation of *CSMD1* expression on gel invasion. Three trans-well chambers of 2 representative clones and parent cells are displayed (for full dataset see S7). The invasion index of generated clones vs. parent is depicted as bar charts (white and black bars respectively). Left: *CSMD1*-expressing clones vs. *CSMD1*-negative H103 parent cells. *CSMD1* expression results in a marked decrease of gel invasion (p=5.975×10^−5^). Right: *CSMD1*-silenced clones vs. *CSMD1*-expressing BICR16 parent cells. *CSMD1* silencing results in a marked increase in gel invasion (p=1.822×10^−05^).

### Later changes in evolution of HNSCC

SCNAs were analysed in two panels of tumour cell lines (S10 Fig) using two different approaches (SNP array and array CGH). Since there was little difference between the two panels in both high copy number alterations (gains >2 and homozygous deletions) and low copy number alterations (gains ≤2 and hemizygous deletions), the data were merged for further analyses.

#### Progression to HNSCC is characterised by increased frequency of SCNAs of chromosomal regions involved in PPOLs as well further SCNAs of specific additional regions

Overall, progression to HNSCC was characterised by an increased frequency in the SCNAs observed in immortal PPOLs (Fig. 2) with statistically significant increases in losses of proximal part of Chr3p, Chr4q and Chr18q and gains of Chr20q. In addition, there was additional loss of Chr10p coupled with gains of Chr5p, Chr9q, Chr14q and Chr11q. Some genes (e.g., *CDKN2A, CDKN2B*) showing homozygous deletions in PPOLs showed an increase in frequency of the same in HNSCCs. There were in addition, homozygous deletions of further genes such as *PTPRD*, *LRP1B* and *LINGO2* some of which showed high frequency hemizygous losses in PPOLs. High copy number gains, which were rare in PPOLs, were more frequent and principally centred on chromosome 11q. As with PPOL-derived cell lines significant peaks of copy number gains and losses were identified using GISTIC (26). The genes in the peak regions at different levels of stringency (Q=0.05-0.25) are shown in S6 Tables.

#### Identification of key genes in progression from PPOLs to HNSCC by integrative analyses

In addition to looking at differences in frequency of SCNAs between and early and late lesions in HNSCC progression, we used integrative analyses to further delineate key genes in transition from PPOLS to HNSCC. Gene expression array data were available for 29 samples (11). We identified genes that showed significant correlation of copy number with gene expression, from genomic regions that showed statistically significant difference in frequency of SCNA/LOH between PPOLs and HNSCC cell lines (S11 Table). This identified 67 genes (Fig. 5) of which 50 have been previously reported to be show some association with cancer (including *NOTCH1* and *PIK3CA*) using PUBMED search. Nine of the 50 genes have been previously associated with HNSCC including *DVL3*, and 5 genes (*NOTCH1*, *PPP6C*, *RAC1*, *EIF4G1*, *PIK3CA*) were identified as mutational cancer drivers in IntOGen (Release 2014.12).

**Fig 5.**
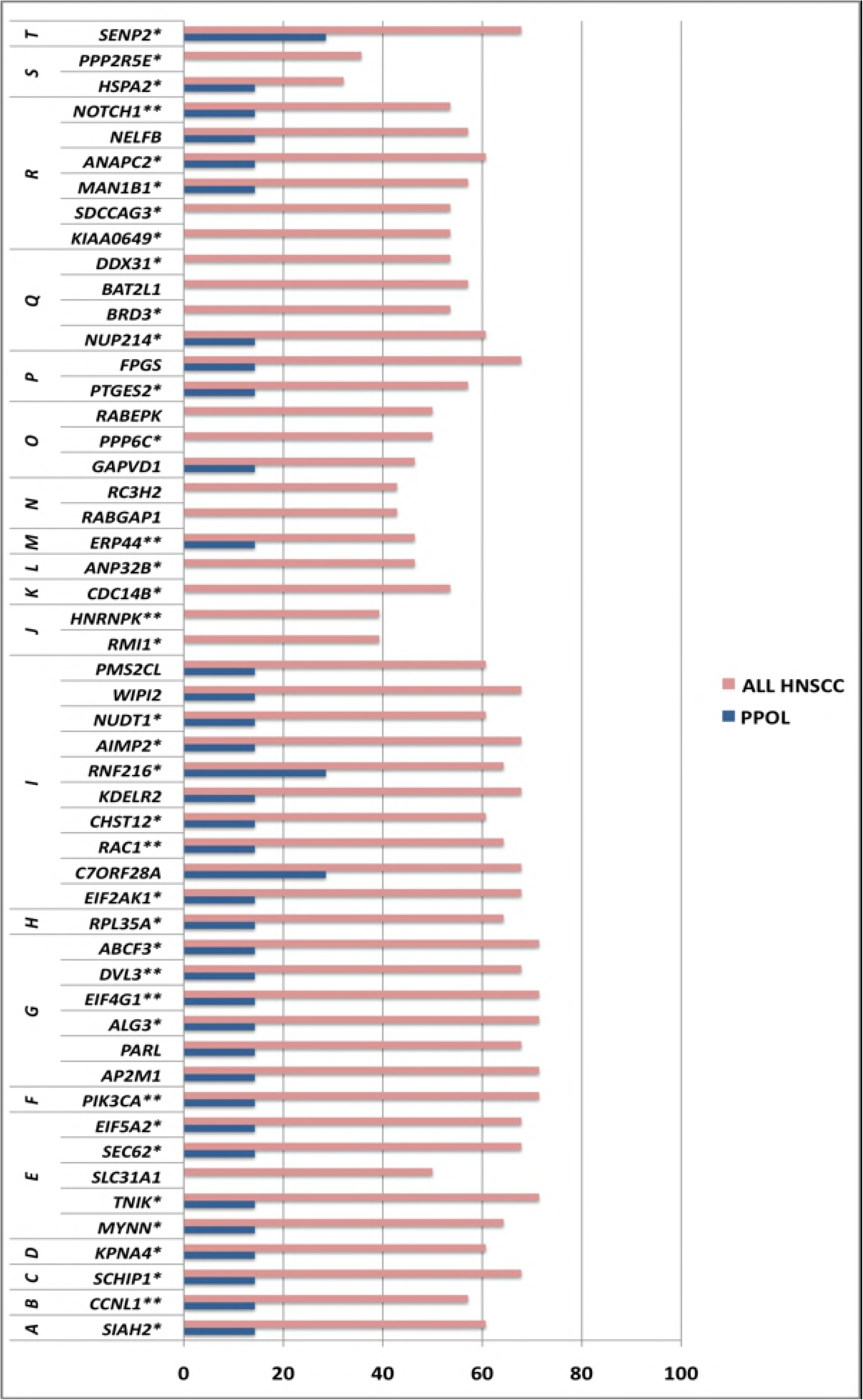
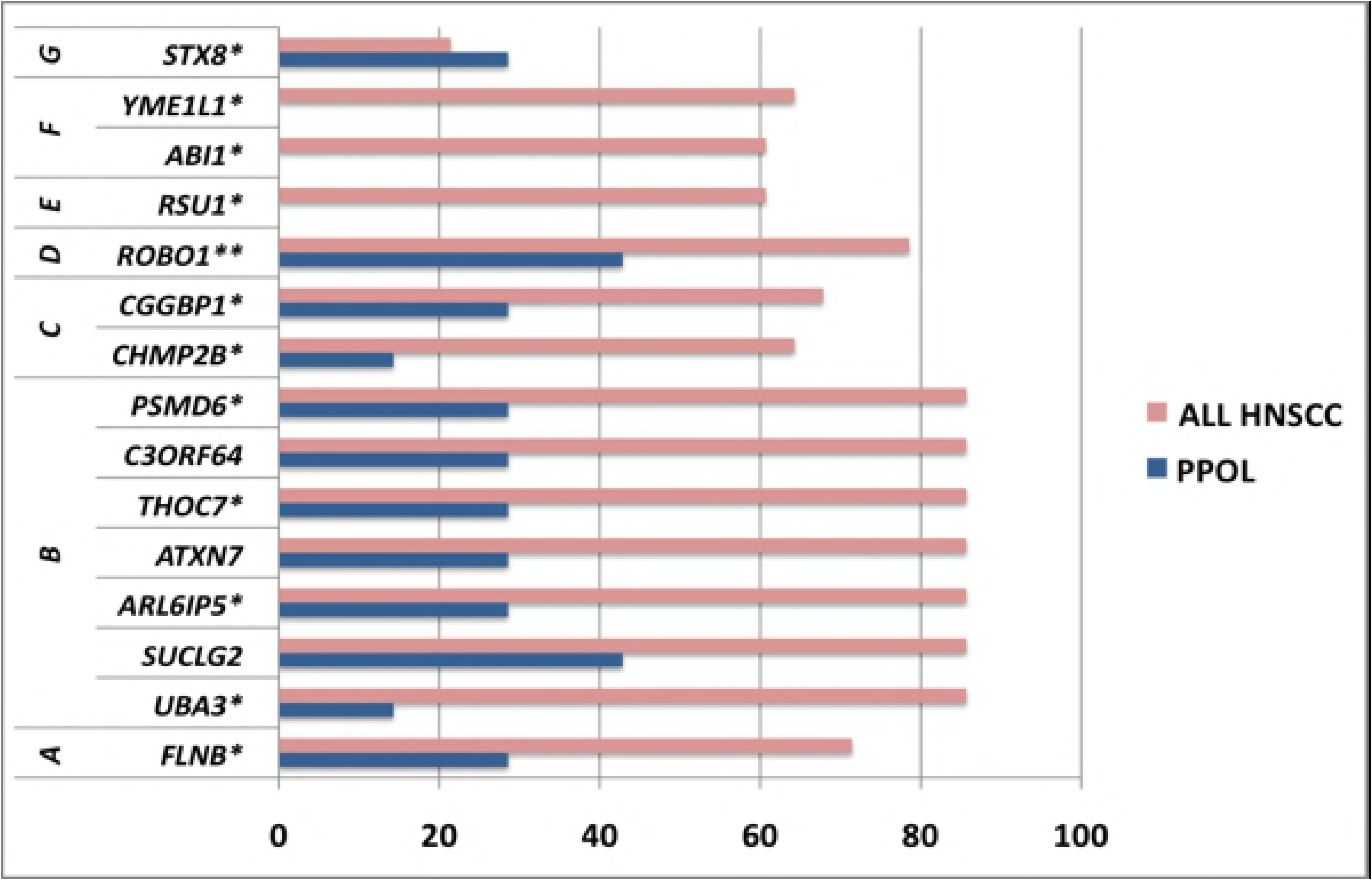
Frequency of SCNA of genes from chromosomal regions showing significant difference in SCNAs between PPOLs and HNSCC and significant correlation with expression. **A.** Copy number gains **B.** Copy number losses. Genes reported to be associated with any cancer previously by PUBMED search (‘Gene name, Cancer’). **Genes reported to be associated with HNSCC previously by PUBMED search (‘Gene name, HNSCC’, oral cancer). Chromosomal regions showing differences between PPOLS and HNSCC are indicated by letters on the left; in top panel (A), the regions are: A, chr3:151842842-152984767; B, chr3:157160169-158933894; C, chr3:160599782-161161335; D, chr3:161705726-162566403; E, chr3:170691970-173911476; F, chr3:177707736-180451354; G, chr3:185035692-185529164; H, chr3:198578155-199298372; I, chr7:1631815-7317208; J, chr9:85367221-85868737; K, chr9:97596715-98774190; L, chr9:99427940-100373274; M, chr9:101504820-101852863; N, chr9:123376788-125083831; O, chr9:126825879-127177239; P, chr9:129518474-129927677; Q, chr9:132985780-136636113; R, chr9:137291321-139534231; S, Chr14:62865875-63862093; T, chr3:186575268-187080482. In bottom panel (B), the regions are: A, chr3:57677987-58154068; B, chr3:61422777-73764765: C, chr3:87035206-88461236; D, Chr3:78706269-79206160; E, chr10:16585949-17022250; F, chr10:26833288-28028304; G, chr3:161705726-162566403; chr17:8028078-9207567. The regions are ranked A onwards in order of decreasing statistical significance.

#### Cell lines from HNSCC with and without lymph node metastases show specific differences in SCNAs

The aggregate HNSCC cell line data concealed notable differences between cell lines from tumours with and without lymph nodal metastases (LN+ve and LN-ve respectively). Lymph node metastases status of 17 of 28 HNSCC lines was known (10 LN+ve; 7 LN–ve). The LN+ve cell lines showed higher frequency of high copy (>2) number gains of a 1.76 Mb region at chromosome 11q13.2-q13.3 (p<0.05) that encompasses 10 genes including *CCND1* and the microRNA hsa-miR-548k (Fig. 2). Similarly, a higher frequency of high copy number gains on two regions on chromosome arm 3q was observed involving *NAALADL2* (chromosome band 3q26.32), and *TP63* and *CLDN1* (chromosome band 3q28). Hierarchical clustering of all HNSCC cell lines by high-copy number aberrations also revealed two major groupings defined by the presence or absence of the high-copy number amplicon on chromosome band 11q13.2-q13.4 (p<0.01, q bound <0.1) (S12 Figure). In the group lacking the amplicon, only 2 of 8 cell lines with known nodal staging, were derived from LN+ve cancers and each of these involved a single node less than 2 cm (TNM stage N1). By contrast, in the group with the amplicon, 7 of 9 cell lines were derived from LN+ve cancers and 4 of the 7 tumours were graded as TNM stage N2 or higher. These results are consistent with very recent findings linking the 11q13.3 amplicon with poor survival in HNSCC patients (34). Comparison of all copy number changes also revealed further important differences. LN+ve cell lines had a higher frequency of copy number gains of chromosome 3q, 12q, 14q and 20, together with a higher frequency of copy number losses on distal regions on chromosome 3p and 11q, and chromosomes 4, and 18q (Fig. 2). Some SCNAs (copy number gains in chromosome regions 7p12.2-21.3 and 9q31.3-32) were more frequent in LN-ve cell lines than in LN+ve cell lines.

#### GISTIC peak regions of SCNA in PPOLs and HNSCCs show significant progressive enrichment of genes involved in cancer pathways

Statistically significant (adj. P<0.01) enrichment of genes in KEGG ‘pathways in cancer’ as well as other specific cancer pathways was observed in GISTIC regions in both PPOLs and HNSCC cell lines (Fig. 6 and S13 Table). This enrichment was observed even if the analysis was limited to 1466 genes in GISTIC regions that showed significant correlation with expression after correction for multiple testing (adj. p<0.05) in immortal PPOL and HNSCC cell lines for which expression data was available. Surprisingly, many genes in GISTIC regions that showed high frequency of SCNAs in HNSCC including known HNSCC cancer drivers (e.g., *CDKN2A*, *MYC)* did not show correlation with gene expression. Therefore, we tested whether integrative analysis was a reliable method to predict *<u>in vivo</u>* protein expression. We examined expression of protein by immunohistochemistry of two genes (*BCL2L1* encoding an apoptosis regulator and *CLDN1* encoding a component of tight junctions in epithelia) that show copy number gains. *BCL2L1* gene showed high frequency/low copy number gains in PPOLs (71%) and LN+ve HNSCC (90%) but not in LN-ve HNSCC (28%). *CLDN1* showed low frequency/low copy number gains in PPOLs (14%) and high frequency/low copy number gains in HNSCC (73% overall, 53% LN-ve HNSCC, 90% LN+ve HNSCCs). *BCL2L1* but not *CLDN1* shows significant correlation of copy number with gene expression in this study (adjP=0.01 and 0.74 respectively). However, in PPOLs and HNSCC biopsies (S14 Fig) both *CLDN1* and *BCL2L1* showed significantly increased expression (p<0.0001) in HNSCC compared to normal tissues and PPOL. This indicated that correlation of SCNA with transcript expression in integrative analyses may not be a reliable surrogate indicator of functional importance of a gene, and that protein expression may better reflect the underlying SCNA.

**Fig 6.**
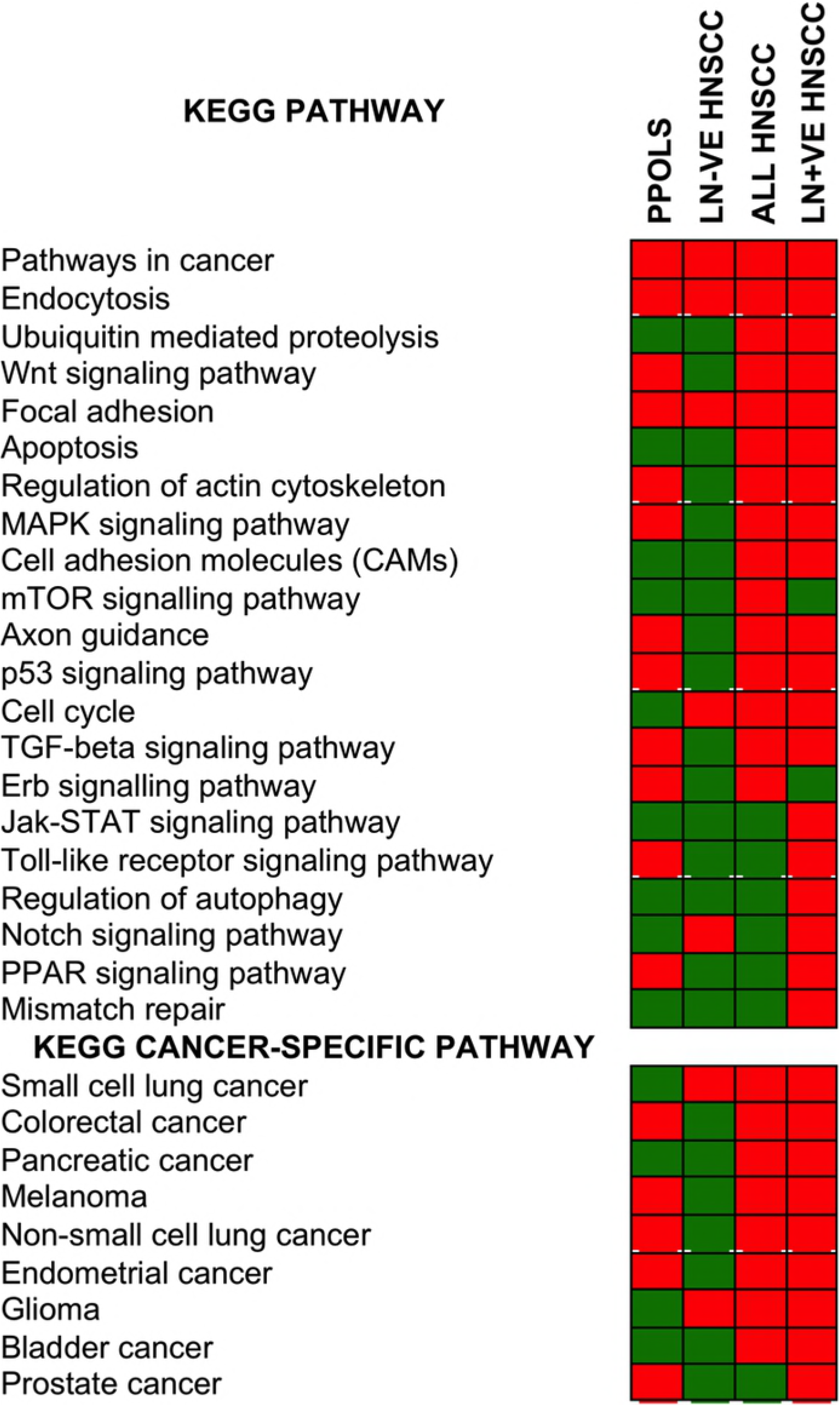
Enrichment of genes in cancer–relevant pathways in GISTIC extended regions in PPOLs and HNSCCs. Statistically significant enrichment (adjusted P<0.01) for specific pathways is indicated by a red box and lack of enrichment by a green box. The data are shown in detail in Supplementary Data 9. The ranges for adjusted P values corrected for multiple testing were: ‘All HNSCC’, 1.97E-12 - 0.0098; ‘LN+ve HNSCC’, 5.76E-14 - 0.0083; ‘LN-ve HNSCC’, 7.52E-06-0.0074; PPOL, 5.34E-09 −0.0007. In each section, the pathways were ranked from top to bottom in order of level of significance in the ‘All HNSCC’ group with highest level of significance at the top.

The increase in both the size and the number of the SCNA regions in HNSCC compared to PPOLs, and the differences in SCNAs between LN+ve and LN-ve HNSCC cell lines, were reflected in the progressive increase and/or differences in the enrichment for relevant KEGG pathways genes (Fig. 6). Given that individual proteins participate in multiple pathways and processes, many of the same genes mapped to multiple cancer-related and other pathways. Some of the pathways identified such as TP53 signalling and the cell cycle, have been well characterised in HNSCC, but others such as axon guidance, actin cytoskeleton and motility, and ubiquitin–proteosome pathway less so.

The examination in our cell line panel, of the frequencies of SCNAs involving genes in individual pathways, allowed us to map multiple pathway changes to stages in HNSCC progression (Fig. 7 and S15 Fig). For example, the peak regions of SCNAs in LN+ve cell lines showed enrichment of specific genes in the TGFB pathway that would predict dysfunction of this signalling pathway (Fig. 7a). High frequency losses of receptors (*TGFBRII*, *ACVR2B* and *BMPR1B),* common *SMAD4,* R-*SMAD2* and inhibitory *SMAD7* as well as other intracellular effectors such as *PPP2R1B* and *RHOA*, were observed. This was coupled with high frequency gains of *CHRD* and *TGFB3*. In addition, there were SCNAs of downstream targets with losses of *CDKN2B* (normally induced by the pathway) and gains of *MYC* (normally down-regulated by TGFB signalling). Similarly, enrichment in SCNAs of genes in the *NOTCH1* pathway was observed in both LN-ve and LN+ve cell lines (Figure 6b). However, there were differences in the frequencies of SCNAs of the individual genes (Figure 6b) in the pathway suggesting different alterations of the pathway. LN+ve cell lines showed almost universal amplification of *HES1* and *DVL3* and universal loss of *KAT2B*. LN+ve cell lines also showed a relatively low frequency (20%) of high and low copy number gains of *NOTCH1*. Gain of *DVL3* and loss of *KAT2B* together with a relatively high frequency of gain of *NUMB* and loss of *MAML2* may be expected to disrupt *NOTCH1* signalling but high frequency gains of the downstream targets *HES1* and *HEY1* may negate these changes or alternatively suggest complex amplification and inhibition of subsets of NOTCH signalling elements. Clearly, these genes act in multiple pathways and it is difficult to determine the effect of gain or loss of any single gene without functional analyses. Thus, copy number gains of *DVL3* may be of significance in WNT signalling pathway or in the cross-talk between the two pathways. Amplification of *HES1* and *DVL3* and loss of *KAT2B* were also observed in LN-ve cell lines as well as PPOLs albeit at a lower frequency. Instead, the LN-ve cell lines were characterised by a high frequency, low copy number gains of *NOTCH1* and *LFNG.* In our small sample set, cell lines with mutations and amplifications of the *NOTCH1* locus were mutually exclusive. Some genes such as *MAML1* and *MAML2* showed both copy number gains and losses possibly indicating further heterogeneity in NOTCH pathway aberration.

**Fig 7.**
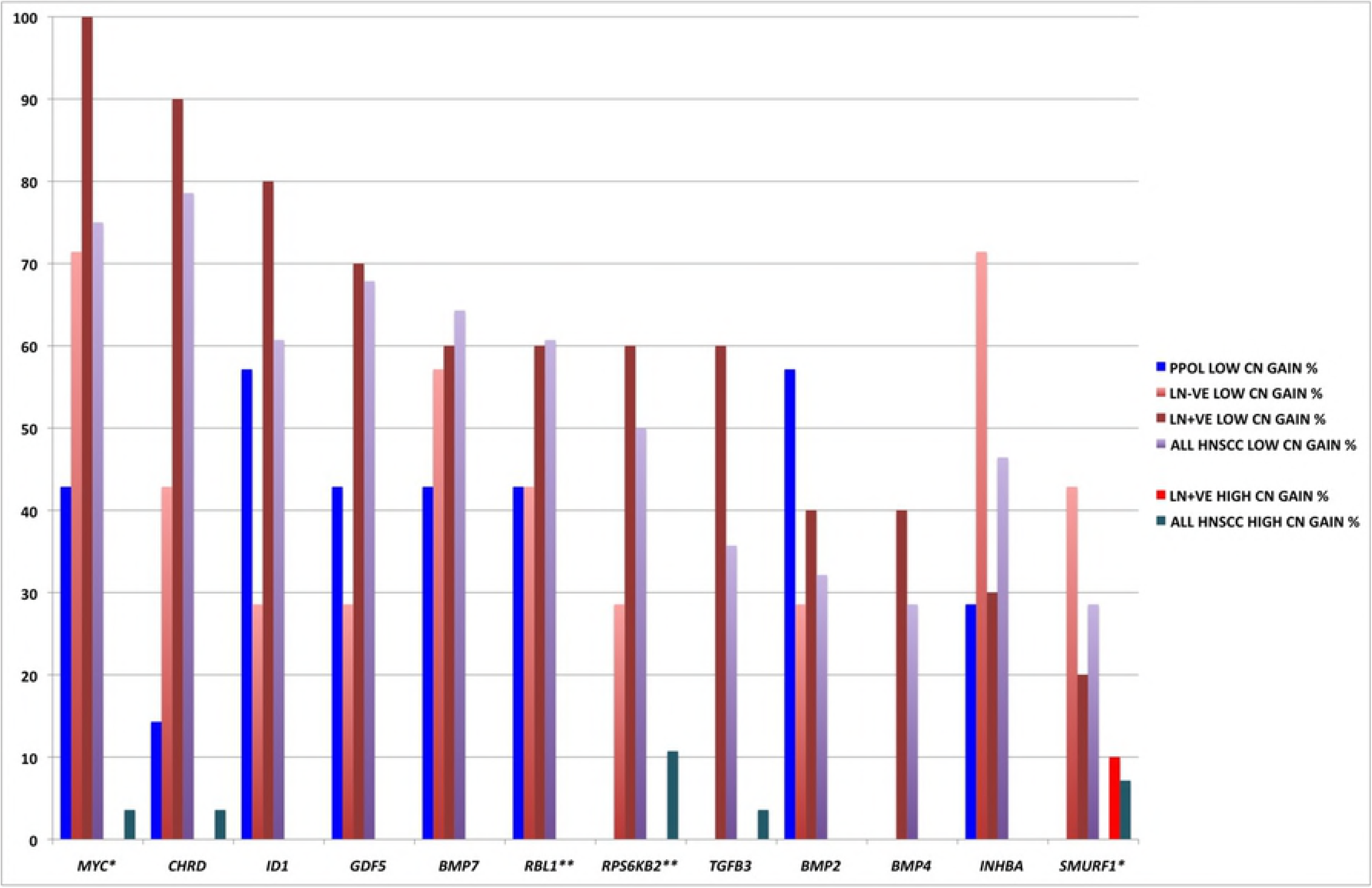
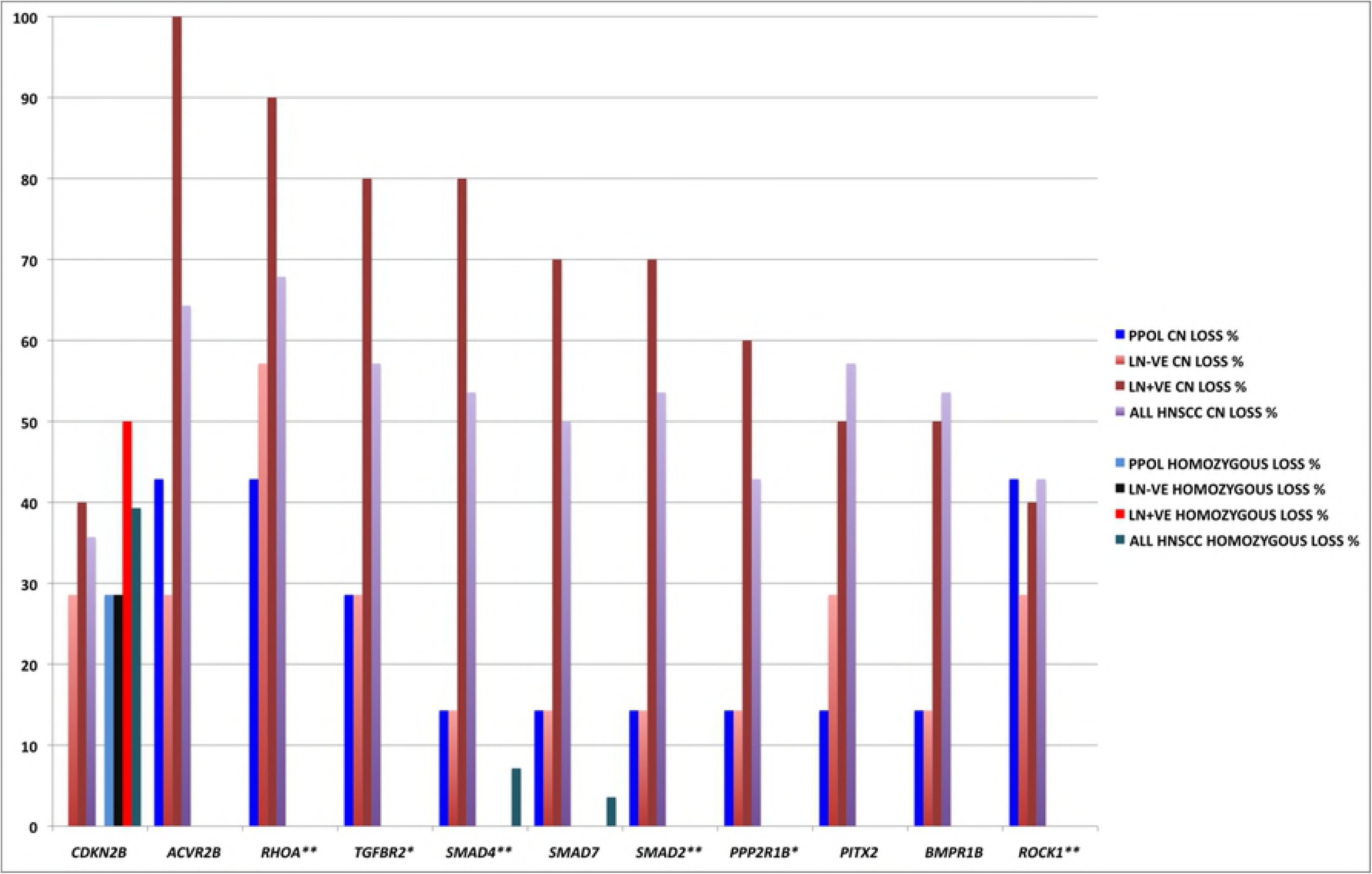
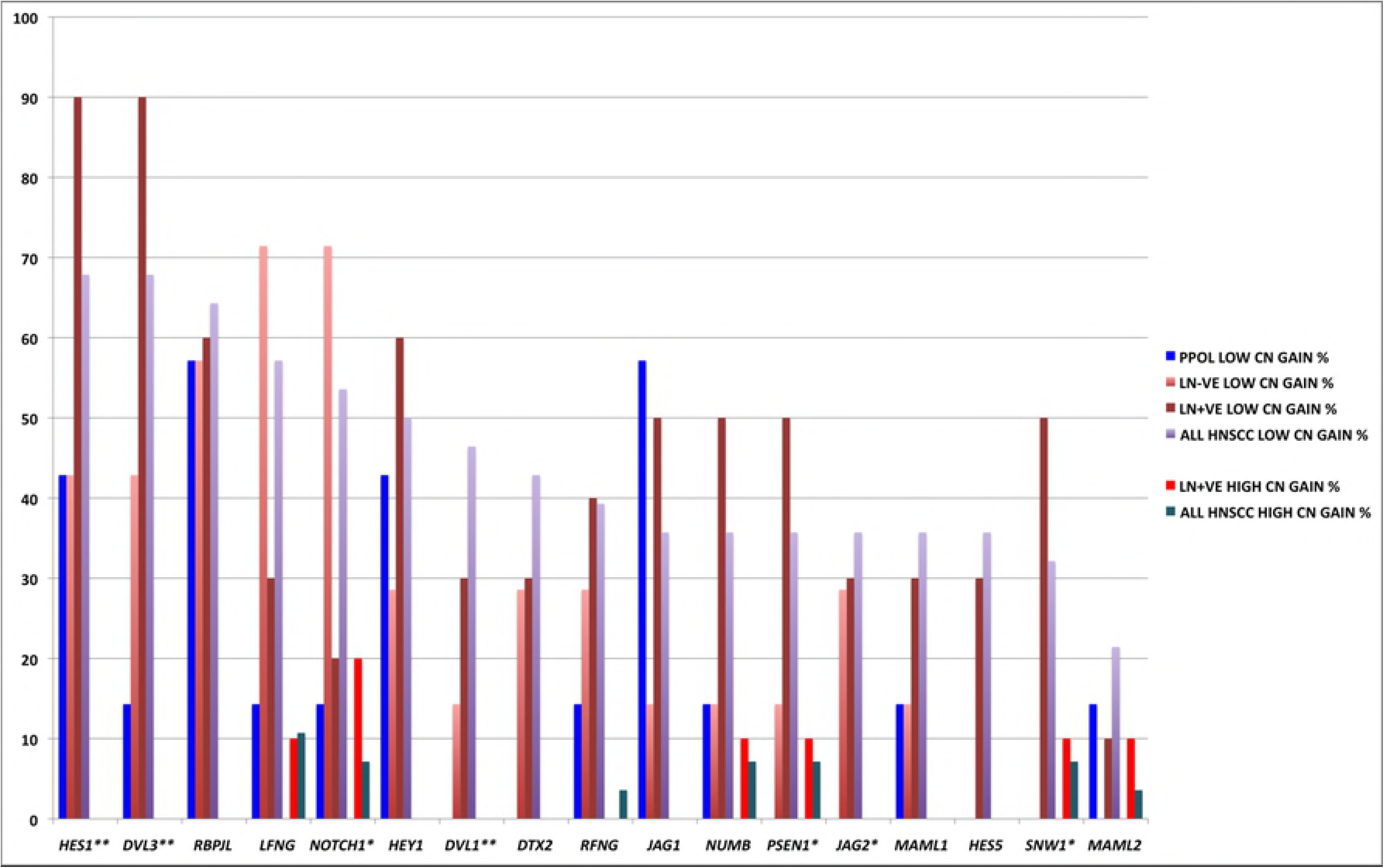
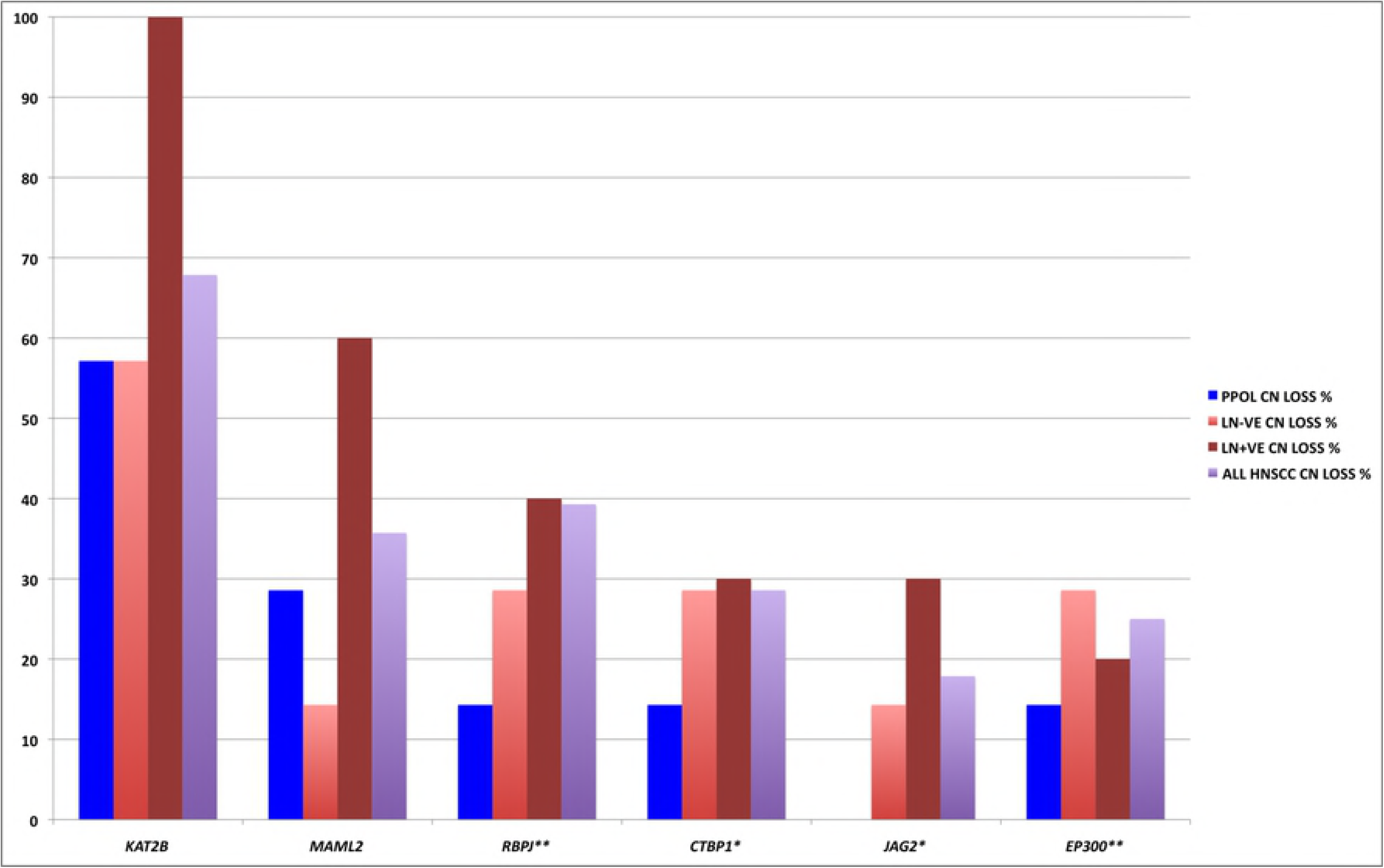
Frequency of SCNAs of genes involved in selected cancer pathways that are significantly enriched in the GISTIC regions in PPOLs and HNSCC. **A.** TGFB pathway **B.** NOTCH pathway. For each pathway, two charts are shown illustrating the frequency of copy number gains (top panel) and losses (bottom panel) in PPOLs, all HNSCC, and HNSCCs with and without nodal metastases (LN+ve and LN-ve respectively). **Genes showing significant correlation with expression in integrative analyses after correction for multiple testing (adj. p<0.05). *Genes showing nominal significance (p<0.05) only are indicated by a single asterisk. Only genes showing at least 40% frequency of SCNA in at least one subgroup, are shown.

#### Multiple genes in SCNA regions may play a role in cancer progression

The prevalence and commonality of deletions and amplifications involving specific chromosomal arms and regions in a wide range of cancers coupled with the enrichment of known cancer-related genes in peak regions of SCNAs and supporting functional evidence for many of the genes in previous studies indicates selection for the SCNAs in cancer cells is driven by the presence of several relevant genes on these chromosomal arms/regions. Clearly, however, pathway analyses will not identify genes such as *CSMD1* which may be cancer-relevant but whose functions are not characterised or mapped to known pathways. We tested this further by examining *ADAMTS9,* a gene not identified in pathway analyses but showing frequent SCNAs.

*ADAMTS9* along with several other genes in this region at 3p14.2, shows a single copy loss in just over 80% of the HNSCC and 29% of PPOL lines suggesting possible further selection of deletions of this or neighbouring genes in cancer progression. *ADAMTS9* is inactivated by promoter hypermethylation in other tumour types including nasopharyngeal carcinoma and oesophageal squamous cell carcinoma (35), (36), Functional analyses suggest a role for *ADAMTS9* in inhibiting angiogenesis (37). In our array CGH data of 347 tumours of multiple sites (unpublished), *ADAMTS9* showed copy number losses in 23% of the tumours (S16A Table) and in Tumorscape (Release 1.6) analyses, it was focally deleted in epithelial tumours (Q=8.38E-6, frequency=0.369). In the present study, we analysed mutations reported in COSMIC (v76, cancer.sanger.ac.uk), (38). Of 40 mutations, 3 were nonsense mutations, and 13 of 37 missense mutations (derived principally from lower gastrointestinal tract) were predicted to be deleterious by Polyphen and SIFT analyses (S16B Table). We failed to identify any likely pathological mutations in our exome analysis and it was not part of our HaloPlex sequencing panel.

We analysed *ADAMTS9* promoter methylation by pyrosequencing in a subset of the HNSCC cell lines where DNA from matching normal tissue was available and also in the DNA from set of primary HNSCC with matching normal tissues. Promoter hypermethylation was observed in both immortal HNSCC cell lines (7/17, ~41%) and primary HNSCC (9/20, ~45%) (Fig. 7A-C). By contrast, promoter hypermethylation was less frequent in PPOL cell lines (1/7, ~14%) and was not observed at all in the mortal PPOL or mortal HNSCC cell lines (Fig. 8) mirroring the pattern of *ADAMTS9* SCNAs in PPOL and HNSCC. We also analysed the expression of the *ADAMTS9* transcript in primary HNSCCs and also in other cancers using Tissuescan panel (Origene, Maryland, USA). In primary HNSCCs, 17 of 23 samples (~74%) showed reduced expression compared to the normal tissues (Fig. 8D-E). In other tumour types, reduced expression was seen in carcinomas of the breast, colon, lung and thyroid (Fig. 8F). These findings suggest a possible role for *ADAMTS9* in HNSCC and other cancers. Similarly, several other genes not previously mapped to any KEGG pathway but within GISTIC regions may be significant. For example, *BOP1* on chromosome arm 8q24.3 is in GISTIC focal region of deletions in both LN-ve and LN+ve HNSCC cell lines (S6 Table). PPOL and HNSCC cell lines showed high frequency, low copy number gains (PPOL, 43%; LN-ve HNSCC, 53%; LN+ve HNSCC 90%) and low frequency, high copy number gains (PPOL 0%; LN-ve HNSCC 14%; LN+ve HNSCC 10%) of *BOP1* and SCNA correlated with gene expression (adj p<0.001) in integrative analyses. *BOP1* has recently been shown to be a downstream target of Wnt signalling and promotes cell migration and metastases in colorectal carcinomas (39) and epithelial–mesenchymal transition, migration and invasion in hepatocellular carcinoma (40).

**Fig 8.**
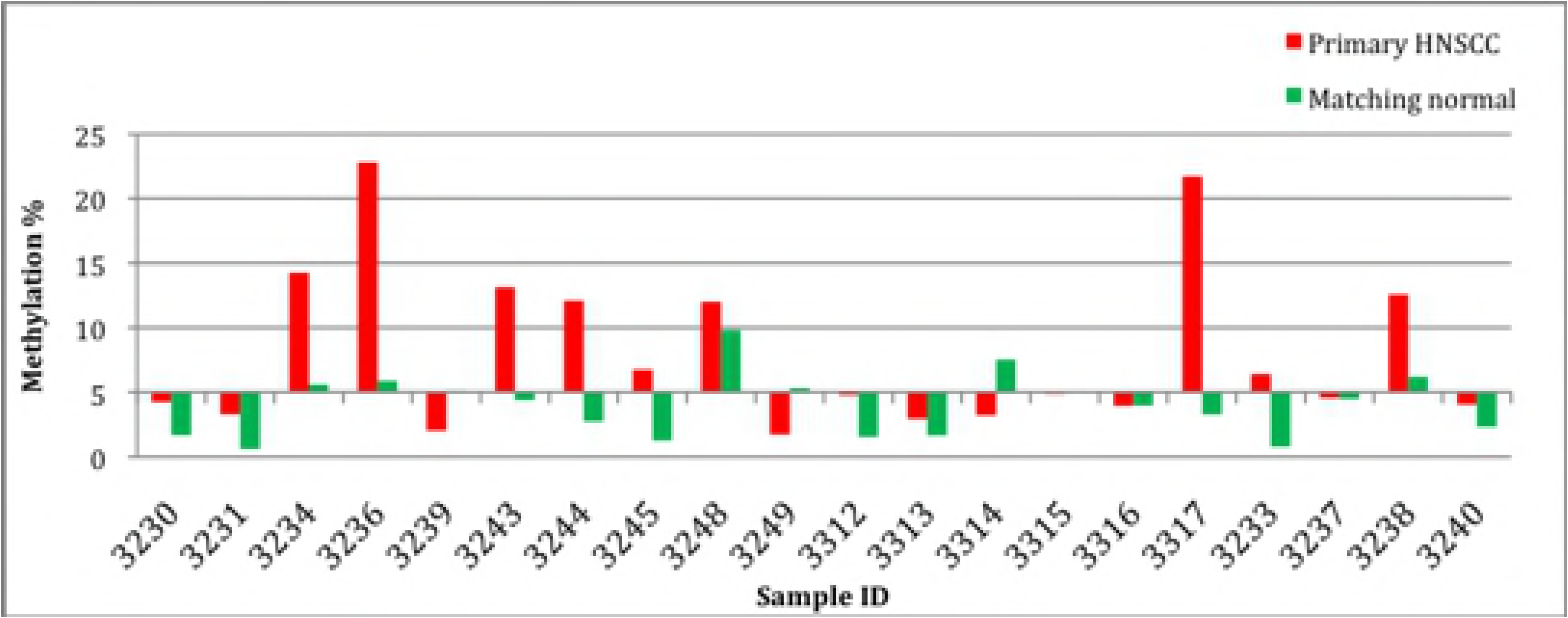
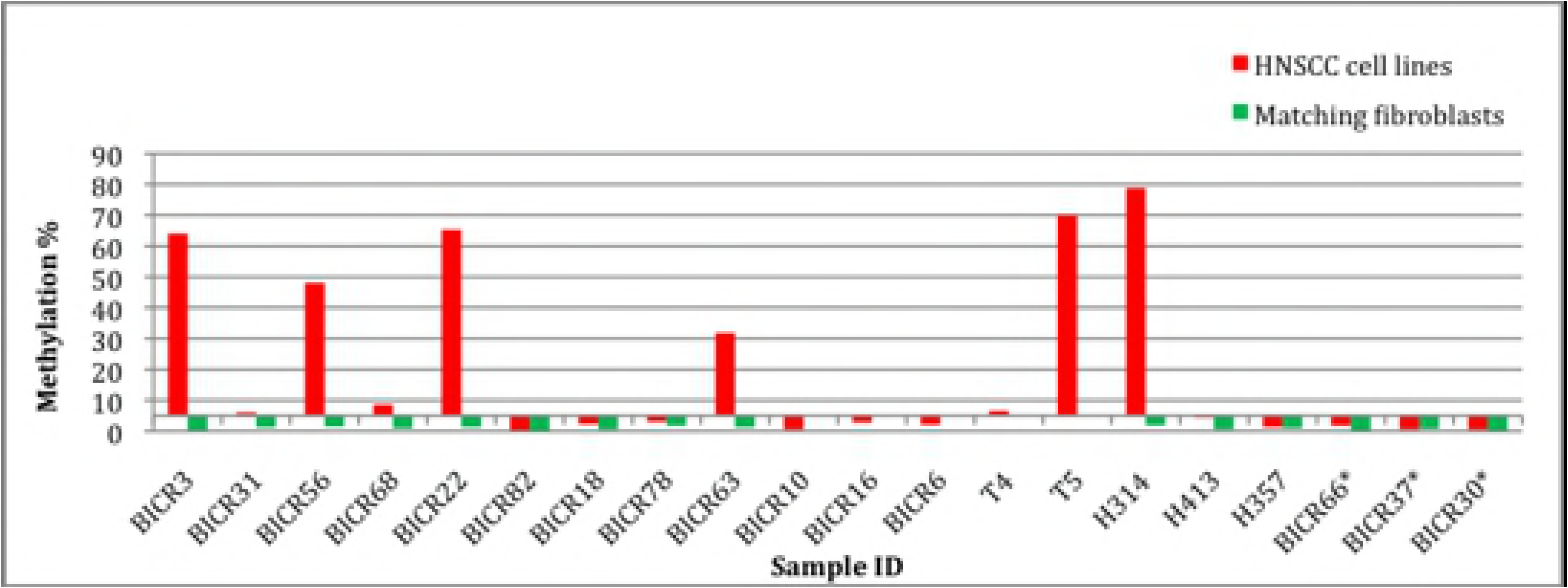
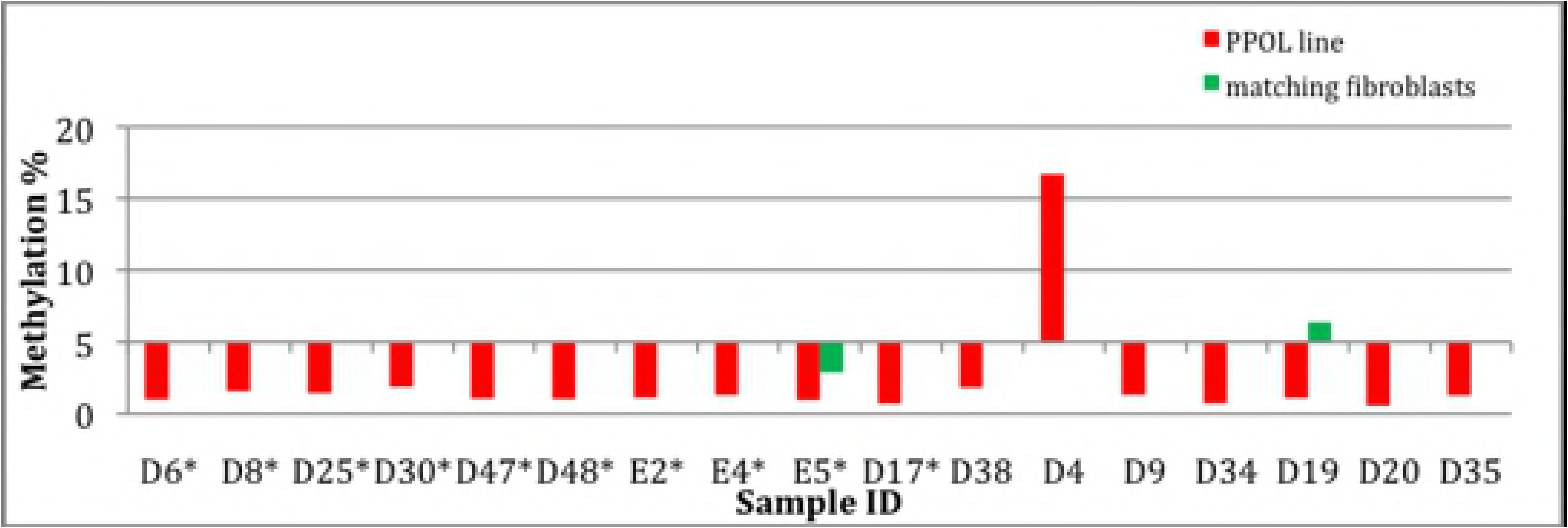
Promoter hypermethylation analyses of *ADAMTS9*. Sample ID is shown on the X-axis and the mean methylation percentage is represented on the Y-axis. For a sample, a mean methylation percentage greater than 5% was considered as significant promoter methylation. **A.** Primary HNSCCs and matching normal samples. Promoter methylation was observed in 9 of 20 (~45%) primary HNSCCs (results from samples 3232, 3241, 3242 and 3247 were excluded from analysis as either the normal or tumour reaction failed). **B.** Mortal and immortal HNSCC cell lines. Promoter methylation was observed in 7 of 17 (~41%) immortal HNSCC cell lines, but not in any of the mortal HNSCC cell lines (indicted by *). **C.** PPOL cell lines/cultures. Promoter methylation was observed in only 1 of 7 (~14%) immortal PPOL cell lines and none of the mortal PPOL cell lines (indicted by *).

## DISCUSSION

Many recent studies have reported genetic and epigenetic changes in HNSCC (4), (3), (5), (6), (7). Here, we have extended these findings by further detailed genomic analyses of a unique panel of cell lines from premalignant lesions (PPOLs) as well as subsets of HNSCCs with and without lymph node metastases. This approach has allowed us to map the genetic changes to the stages of evolution of HNSCCs and to identify the earliest abnormalities, which are significant in tumour progression. The separate analyses of the HNSCC subgroups with respect to nodal metastases has facilitated the identification of changes which may be otherwise masked by looking at average changes in a heterogeneous group.

The prevalence of the changes observed in this study must be regarded with caution given the small sample size and the use of cell lines. Changes observed in cell lines may not be fully representative of primary tumour because of clonal selection and continuing evolution in culture. However, the patterns of changes observed in our study are consistent with studies of primary tumours. Our GISTIC analyses identified similar peak and extended regions as those identified in 3131 primary tumours by Beroukhim et al., 2010 (31). Balanced against this, one major advantage of using cell lines is that extensive heterogeneity of primary tumour samples can mask SCNAs (41), (42), whereas early passage cell lines give cleaner results (43). Nevertheless, it is essential that the findings are verified in larger sample sets of primary premalignant lesions and tumours.

We have also demonstrated that techniques such as GISTIC, pathway and integrative analyses for identifying the pertinent or significant changes amongst the myriad changes observed in tumours have limitations. *BCL2L1* and *CLDN1* both with high frequency copy number gains in HNSCC, show increased protein expression in primary tumours despite lack of correlation in integrative analyses for *CLDN1*. *ADAMTS9*, which is in not in a GISTIC peak region in our sample set and does not map to a KEGG pathway, showed frequent copy number loss and promoter hypermethylation together with decreased gene expression in HNSCC but not in PPOLs or normal tissues indicating potential role in HNSCC.

Given the relatively low frequency (~30% or less) of sequence mutations of genes other than *TP53* both in the current and previous studies, together with prevalence of consistent and frequent SCNA changes across wide range of tumours (31), it is likely that the selection processes in the clonal evolution of these tumours is driven by these SCNAs. Additionally, we show here that the earliest changes are often characterised by focal SCNAs (for example, deletions at chromosome 8p23 involving *CSMD1*) with further selection for loss of whole arm or large region of arm during progression or evolution. This suggests that regions of SCNAs harbour multiple genes that collectively provide selective growth advantage. This is further supported by the KEGG pathway analyses, which show statistically significant enrichment of genes in the cancer relevant pathways in peak regions of SCNAs identified in HNSCC using GISTIC. Additionally, there are numerous functional studies in the literature of different genes in the same chromosomal regions, which are involved in development of tumours of same type. Although this suggests that there are a large number of cancer drivers, it is possible that the genes do not represent primary drivers but their alterations provide a further cumulative selective advantage against a background of primary driver loss-of-function or gain-of-function.

In the present study, we confirm previous observations (23), (24), (16), (25) that loss-of-function of *TP53* (primarily through sequence mutations) and *CDKN2A* (through SCNA, promoter hypermethylation and sequence mutations) are early changes in HNSCC development. Furthermore, we demonstrate for the first time that loss-of-function sequence mutations in *NOTCH1*, *KMTD2* (*MLL2*) and *FBXW7* are present in PPOLs and represent early but less common changes.

In accordance with previous observations (8), we demonstrate that a loss of chromosome arms 3p, 8p and 9p and gains of chromosome 20 are the earliest changes in HNSCC Using cell lines from progressive and non-progressive PPOLs, we show that the earliest changes are characterised by focal deletions and/or promoter hypermethylation of *CSMD1* on chromosome arm 8p23.2. The role of *CSMD1* in cancer is relatively unknown but deletions at this locus have been reported in many types of cancers (31). Like many other commonly homozygously deleted genes in HNSCC and other tumours, *CSMD1* (and *FHIT*) and are large genes that are located in regions of low gene density or ‘gene deserts’ (S17 Table). Deletions appear to occur more frequently in such regions and thus, at least some of these SCNAs may be aetiologically unrelated to cancer development and over represented through low selection pressure against these changes (31). Furthermore, in this study, a few pseudogenes in ‘gene deserts’ were also found to have sustained homozygous deletions (Supplementary Data 17) supporting the notion that at least of some of these SCNAs are ‘non-specific’ changes. Nevertheless, there is convincing functional evidence supporting a tumour suppressor role for at least some of these genes including *FHIT* (30), (29) and other large genes such as *DCC* that are located in gene deserts (44). In the present study, we provide functional evidence for the first time for a tumour suppressor role for *CSMD1* in head and neck squamous mucosa. Our findings support findings in breast cancer reported by (45). *CSMD1* encodes for a predicted transmembrane protein with a multidomain extracellular structure that is likely to act as a multi-ligand receptor mediating endocytosis of ligands. However, this remains to be characterised. One previous study has reported functional analyses in melanoma cell lines demonstrating a role for *CSMD1* in reducing proliferation and invasive potential possibly through *SMAD* pathway (46).

In this study, whilst we did not detect mutations in *FAT1* (a gene known to be mutated in HNSCC) in PPOL, we did identify hemizygous and homozygous deletions of the gene confirming this gene as an early target for inactivation in HNSCC development. In addition to *FAT1,* we also identified novel homozygous deletions in *NCKAP5*, *SORBS2* and *FAM190A* in PPOLs. *SORBS2* has been shown to induce cellular senescence (47). *FAM190A* is a structural or regulatory component of mitosis and its loss may contribute to chromosomal instability (48). Little is known of *NCKAP5* and we demonstrated that this is not a frequent target of SCNAs or reduced expression in HNSCC or other tumour types. However, its product interacts with product of *NCK1*, a gene in the extended GISTIC region on chromosome arm 3q; frequent amplifications of *NCK1* in LN+ve HNSCC cell lines used in this study and pan-tumour low frequency gain-of-function mutations in the IntOGen dataset suggest a possible novel cancer-relevant pathway. Interestingly, NCK proteins are essential signalling elements in cytoskeleton organisation and cellular motility (49) and in the present study, there was a significant enrichment of the genes in the cytoskeleton organisation and cell motility pathways in the extended GISTIC regions in this study. NCK1 has been reported to be necessary for EGFR-mediated migration and metastases in pancreatic cancer (50) and depletion of NCK1 increases UV-induced TP53 phosphorylation and apoptosis (51).

In contrast to study by Bhattacharya and colleagues (8), this study did not find evidence of significant subset of immortal PPOLs or HNSCCs that didn’t show loss at 8pter-p23.1 together with gains on 3q24-qter, 8q12-q24.2, and chromosome 20. However, our sample size is relatively small and there may be a selection in culture of the cells with these genetic alterations. Interestingly, in the previous study (8) the subgroup lacking these genetic alterations showed genetic stability, lack of *TP53* mutations and a much-reduced predisposition to metastasis. It is interesting to speculate whether these correspond to ‘mortal’ PPOLs and HNSCC cultures, which are genetically stable and also lack *TP53* mutations.

Our findings with respect to mutations and SCNA are similar but not identical to those reported by (18) in synchronous dysplasia and HNSCC. In our study, we were able to show progressive changes with transformation to malignancy and lymphovascular spread. We note that in their study only a minority of low-grade dysplasias showed changes that were present in high-grade dysplasia and HNSCC, and they suggested that SCNAs were not necessary for the low-grade dysplasias to develop. We interpret this with caution as unsurprisingly, low-grade dysplasia can be very difficult to distinguish from normal tissues with absolute certainty and agreement between histopathologists is generally weak in grading low-grade dysplasia. Thus, some low-grade dysplasia lesions may represent genuine dysplasia whilst others may present normal tissues. Regardless of this, what is clear from our study is that the SCNAs arise after the breakdown of cellular senescence. The mortal PPOLs do not display SCNAs. Additionally, the loss of expression of *CDKN2A* is nearly ubiquitous in our PPOL panel (16) and in PPOL tissues *in vivo* (24),(52), (11). Furthermore, PPOL D17, which has lost CDKN2A expression whilst remaining mortal and retaining normal *TP53* and telomerase status, has only minimal chromosomal gains and no losses, suggesting that CNAs follow breakdown of senescence.

The results of this study demonstrated that GISTIC extended SCNA regions in the PPOLs and in HNSCC with and without nodal metastases, harbour genes that are implicated in a number of cancer-relevant KEGG pathways. Several of the genes that we identified have been reported previously as being associated with HNSCC but the present study is the first to systematically identify these genes and others, in the context of cancer-relevant pathways and map them to cancer progression. We have described this in context of TGFB and NOTCH signalling pathways and provided supplementary data for other cancer–related pathways.

Our data show enrichment of genes of the TGFB signalling pathway in the GISTIC regions and higher frequency of copy number loss associated in LN+ve HNSCC compared to LN-ve HNSCC and PPOLs. Disruption of TGFB pathway in HNSCC is well established (53). *Smad2*-null mice, for example, develop spontaneous HNSCC and consistent with our findings in this study, copy number losses of *SMAD2, SMAD4* and *TGFBRII* are associated with an aggressive tumour phenotype and lymph node metastases (54), (53). In the present study, we also report relatively common SCNAs of other genes in TGFB pathway that have seldom been reported and or not at all in HNSCC. These anomalies include copy number losses of activin/BMP receptor *ACVR2B,* downstream and SMAD-independent pathway effector *RHOA*, together with copy number gains of BMP inhibitor chordin (*CHRD*) and *ID1*. ID1 is a helix-loop-helix protein that has been shown to induce immortalization in keratinocytes (55) and overexpression of the protein has been reported recently in HNSCC (56).

*NOTCH1* mutations (4), (3) and *NOTCH1* pathway alterations have been reported in HNSCC (5). We have extended these findings and demonstrate that *NOTCH1* mutations are present in two of the three progressive PPOLs but not in any of the non-progressive lesions. This indicates that *NOTCH1* inactivation is a relatively early event but still consistent with previous observations that suggest that *NOTCH1* inactivation plays a key role in progression to invasive carcinoma of already initiated cells (57). Our data are also consistent with that of Agrawal and colleagues (4) who have shown no association or mutual exclusivity of *NOTCH1* and *TP53* mutations. Interestingly, the results of the present study demonstrate that SCNAs of several genes in the NOTCH1 signalling pathway including *NOTCH1*, are more frequent then loss-of-function mutations of *NOTCH1*. Furthermore, many (but not all) of these SCNAs are gain of function changes in the NOTCH1 pathway. This is consistent with recent observations in primary HNSCC demonstrating over-expression of both ligands and receptors in this pathway (58), (59). Although mainly loss-of-function mutations in *NOTCH1* have been reported in HNSCC to date, activating mutations in HNSCC in a Chinese population have also been described recently (60). Our data, therefore add support to the emerging consensus for dual oncogenic and tumour suppressive role for *NOTCH1* in HNSCC although further functional analyses are necessary to confirm this proposal. Some SCNA’s of *NOTCH1* pathway genes such as copy number gains of *DVL3* may also be significant in the context of cross-talk with the WNT signalling pathway.

In this study, we identified a potentially deleterious *NOTCH1* sequence variant in two immortal poorly differentiated PPOL lines and in two immortal and one mortal HNSCC lines. If the mutation in the mortal line which has a wild type p53 (57) and no detectable CNVs (this study) or LOH (15) is pathological as predicted, it would suggest that *NOTCH1* mutations are independent of genomic instability. The presence of *NOTCH1* mutations in this line also offers a plausible explanation for the poor differentiation of this line in both suspension (15, 61) and surface culture (15) and is also consistent with recent data showing that the knockdown of *NOTCH1* expression in human keratinocytes recreates a poorly differentiated epithelium reminiscent of dysplasia (62). Furthermore, *NOTCH1* has been shown to mediate keratinocyte stratification (61, 63) and stem cell maintenance (64). However, *NOTCH1* deletion has also been shown to promote tumourigenesis and tumour progression through paracrine effects (65), the former of which would also be consistent with *NOTCH1* mutations in PPOLs. Moreover, this last observation would be consistent with the reports of *NOTCH2* and *NOTCH3* mutations in human SCC (3) because loss-of-function of these paralogues promotes tumourigenesis in mouse skin in a paracrine fashion but does not replicate the effect of *NOTCH1* deletion on keratinocyte differentiation (65). It has also been reported that *NOTCH1* is a TP53 target gene (61) and as most of the immortal PPOL and HNSCC lines have TP53 mutations this could be an additional mechanism of its inactivation and is consistent with the altered regulation of hairy enhancer of split 2 (*HES2*) in these lines (11, 61).

Integrative analyses in a subset of samples for genes from genomic regions showing a significant difference in frequency of SCNA between PPOLs and HNSCC identified 67 genes that showed correlation with gene expression including *NOTCH1, PPP6C, RAC1, EIF4G1*, *PIK3CA* and *DVL3*. The role of *NOTCH1* and *PIK3CA* in HNSCC is well established. *PPP6C, RAC1* and *EIF4G1* are also likely cancer drivers according to IntoGen. *RAC1* activation has been reported previously in HNSCC and mediates invasive properties of HNSCC (66); our observation of copy number gains in tumour progression is consistent with this finding. *EIF4G1* is a translation initiation factor that is part of the multi-subunit complex EIF4F that facilitates recruitment of mRNA to the ribosome. This is a rate-limiting step protein synthesis initiation phase. *EIF4G1* is amplified in many tumour types in TCGA data sets (67). Over-expression of *EIF4G1* promotes tumour cell survival and formation of tumour emboli through increased translation of specific mRNAs in inflammatory breast cancer (68). Components of EIF4F are also targets of C-MYC and initiate further translation of specific targets including C-MYC (69). *PPP6C* loss-of-function mutations have been reported in melanomas (70) but in the present study, we observed copy number gains particularly in LN-ve HNSCC; the significance of this observation is unclear. *DVL3* is a transducer of both canonical and non-canonical Wnt signaling pathways (71). Association with HNSCC has not been reported previously but it is amplified in many tumour types in TCGA data sets (67) and inhibition of Wnt signalling in HNSCC results in inhibition of growth and metastases (72).

In conclusion, we have further characterised specific genetic changes that mark progression in head and neck squamous cell carcinogenesis. Although genomic landscapes and progression models of SCCHN (14), (4), (6), (5), (3) have been published previously, we have been able to use our well characterised cell line panels to tentatively assign genetic changes, including novel ones, to specific stages of progression in transcriptionally distinct mortal and immortal classes (9), (11) of the disease and also to cell function.

## Materials and Methods

### Samples

This study was approved by the UK National Research Ethics Service Research Ethics Committee (08/H1006/21).

Details of the samples are shown in Supplementary Data 1. For SNP array, the sample set consisted of 16 HNSCC cell lines, 7 PPOL cell lines and 11 mortal cell cultures derived from PPOL. DNA from matching fibroblasts was available for 6 HNSCC cell lines, 1 immortal PPOL cell line and 2 mortal PPOL cultures. The sample culture conditions were as described previously (15) and DNA was prepared from cell lines using standard protocols. The sample set for array CGH consisted of 12 HNSCC cell lines.

### SNP array analyses

SNP genotyping of the primary HNSCC panel was performed using the Illumina HumanHap550 Genotyping Beadchip and Infinium Assay II as per standard protocols. DNA from the cell lines was quantitated with NanoDrop (Thermo Scientific) and 750ng was used per assay.

### Array CGH

ArrayCGH data were kindly provided by Dr Simon Deardon, AstraZeneca, UK.

### Data analyses

The average genotype call rate was 98.25%; genotype data from two samples with a call rate of <95% in BeadStudio v3.1 (Illumina) were excluded from analyses. Over 75% of samples had GenTrain score (measure of reliability based on the total array of calls for a given SNP) of ≥0.7 and none were below 0.4. Data were pre-processed in GenomeStudio v2009.1 (Illumina) and imported into Nexus Copy Number v5.1 (BioDiscovery, Inc., CA, USA) and OncoSNP v2.7 (73) for further analyses. ArrayCGH data from a second HNSCC panel were also imported into Nexus Copy Number v7.5 (BioDiscovery, Inc., CA, USA).

The robust variance sample QC calculation (a measure of probe to probe variance after major outliers due to copy number breakpoints are removed from the calculation) in Nexus Copy Number v7.5 (BioDiscovery, Inc.), was used to assess the quality of the samples. Data for samples with a score > 0.2 were excluded and the score for the remaining samples was in the range 0.03-0.20. For identification of copy number and copy neutral changes, the BioDiscovery’s SNPRank Segmentation Algorithm was used with significance level of 1 x10^−6^ and a minimum number of probes per segment of 5. Thresholds for determining copy number variation were set at −1 for homozygous deletion, −0.18 for hemizygous deletion, 0.18 for gain (single copy gain) and 0.6 for high gain (2 or more copies). An area was considered to be showing LOH if 95% of the probes in the region had a B allele frequency of >0.8 or <0.2 (homozygous frequency and value thresholds of 0.95 and 0.8 respectively). Allelic imbalance was defined as 95% of the probes in the region showing a B allele frequency of between 0.2 and 0.4 or 0.6 and 0.8 (i.e. heterozygous imbalance threshold of 0.4).

Areas of the genome with a statistically high frequency of aberration (Q-bound value <= 0.5—0.25 and G-score cut-off <=1) after correction for multiple testing using FDR correction (Benjamini & Hochberg), were identified using the GISTIC (Genomic Identification of Significant Targets in Cancer) approach (26).

Group comparisons were made in Nexus with differences in frequency of specific events at any chromosomal location tested for significance by two-tailed Fisher’s Exact Probability Test with an accepted significance level of p<0.01 at a defined level of percentage difference. More stringent Q-bound values based on two-tailed Fisher’s Exact Probability Test corrected for multiple testing using Benjamini-Hochberg FDR correction (Benjamini & Hochberg) as well as a minimum of set percent difference in frequencies between the two groups; significance was accepted at <0.25.

### Promoter methylation analyses

Genomic DNA (approximately 1mg each) from all the HNSCC and PPOL cell lines and matched primary HNSCC samples used were subjected to bisulfite treatment using the EZ DNA methylation™ kit (Zymo Research, U.S.A) according to the manufacturer’s protocol. 30ng each of the bisulfite-treated DNA was used for the pyrosequencing reaction.

PCR and sequencing primers for the pyrosequencing methylation analyses of the CpG rich promoter region were designed using the pyro-Q-CpG software for the genes *ADAMTS9* and *CSMD1.* The forward primer was biotinylated and was used in low concentration (5pmol) along with amplification cycles of 45 to exhaust the primers. The forward and reverse primer sequences used in the study were *ADAMTS9*- F-5’agagatttttaaagttaaaagttgg3’, R-5’tccctcctaccctcctta3’ with the sequencing primer 5’cctcctaccctcctta3’ and *CSMD1*-F-5’ gtagttttagatagatagagtttagttt3’, R-5’ acaaatctcctttctcca3’ with its sequencing primer 5’ aaatctcctttctccaacct3’. Optimized annealing temperature for *ADAMTS9* and *CSMD1* PCR primer pairs is 54°C. Using bisulfite-treated DNA as a template, regions of interest were amplified by standard PCR cycling conditions in a 96-well plate using Qiagen’s Hot start Taq Polymerase to avoid nonspecific amplification. The specificity of the PCR products was then verified by agarose gel electrophoresis. For the pyrosequencing reaction, the PCR product was made single stranded by immobilizing the incorporated biotinylated primer on streptavidin-coated beads. The sequence run and analysis were done on the PyroMark™ Q96 MD pyrosequencer (Qiagen) according to the manufacturer’s instructions. The sequence runs were analysed using the Pyro Q-CpG software. The peak heights observed represented the quantitative proportion of the alleles. The software generated methylation values for each CpG site and also the mean methylation percentage for all the CpG sites analyzed.

### Generation of stable *CSMD1*-expression modulated clones

A panel of nine OSCC cell lines was profiled for *CSMD1* transcript and protein expression status (data not shown). This identified the *CSMD1-*expressing cell line BICR16 and the *CSMD1* non-expressing cell line H103.

BICR16 cells were used to generate stable *CSMD1*-silenced monoclone and polyclone lines with HuSH 29mer pRS shRNA (Origene, Rockville, MD, USA). The degree of silenced *CSMD1* expression level was confirmed by RT-qPCR and flow cytometry assays. Primer sensitivity assays determined the limit of *CSMD1* transcript detection at five ORF copies per light-cycler well.

Stable forced *CSMD1*-expressing monoclone cells were generated by transfection of 15.5kb pCMV6-*CSMD1* expression plasmid (Origene, Rockville, MD, USA) into the *CSMD1*-deleted cell line H103. The plasmid was linearized at the SexAI restriction site within a predetermined 14% non-essential region and used to generate seven *CSMD1*-expressing monoclone cell lines using standard methods. The degree of forced *CSMD1* expression was confirmed by RT-qPCR and flow cytometry assays.

Cell proliferation was determined with CellTiter 96® Aqueous Cell Proliferation MTS Assay (Promega, Southampton, UK) as per the manufacturer’s protocols. Cell growth was calculated as a percentage growth change from the 24-hour time point and population-doubling times were determined. Gel-invasion was assayed using trans-well BD Bio-coat Matrigel Invasion Chambers and control wells (BD Biosciences, Oxford) as per manufacturer’s protocols. Optimal seeding densities were determined empirically. Triplicate Matrigel invasion chambers were used for each clone from a minimum of two different Matrigel batches. Three fields of view were captured for each Matrigel or control chamber (outer, middle, centre areas, each at 120° rotation from one another). Percentage invasion was calculated for each clone and expressed as an invasion index (ratio of clone to parent percentage invasion). Statistical analysis was performed in IBM SPSS 20 & 22 (Wilcoxon Sign-Rank Test) with alpha levels set at 0.05.

### Integrative analysis

Before correlating the SCNA and gene expression values corrections were applied for polyploidy and heterogeneity.

When considered a by-product of instability rather than a response of biological significance, the copy number (CN) values were altered to remove the ubiquitous chromosomal amplification observed in polyploid samples. To do so, all the CN values inferred for SNPs in chromosome arms with mean CN larger than 2.5 were reduced by one unit. The reduction was however rejected in the cases of heterozygous copy neutral calls (CN2 LOH0), copy losses (CN1) and homozygous deletions (CN0). These states were not altered.

Due to heterogeneity, the gene expression values obtained did not represent the expression of the CN-altered cells solely. For each genomic region, a non-negligible proportion of cells do not harbour any alteration. The proportion of cells with normal heterozygous copy number in each region was estimated. When investigating the correlation between SCNAs and mRNA expression the weighted mean CN and LOH values of this mixture were used.

### Exome sequencing

Targeted enrichment and sequencing were performed on 1-3 µg of DNA extracted from the cell lines. Enrichment was performed using the SureSelect Human All Exon 50 MB v4 Kit (Agilent, Santa Clara, CA, USA) for the Illumina system. Sequencing was carried out on a HiSeq 2500 sequencer (Illumina Inc, San Diego, CA, USA), following the manufacturer’s protocols.

### HaloPlex Sequencing

Targeted enrichment and sequencing were performed on 225ng of DNA extracted from the cell lines. Enrichment was performed using a custom HaloPlex Kit (Agilent, Santa Clara, CA, USA) targeting 41 genes. Sequencing was undertaken on a MiSeq sequencer (Illumina Inc, San Diego, CA, USA), following the manufacturer’s protocols.

### Sequence data analysis

Raw paired-end reads were trimmed using Trimmomatic v0.33 to a minimum length of 30 nucleotides. Illumina Truseq adapters were removed in palindrome mode. A minimum Phred quality score of 30 was required for the 3’end. Single end reads as well as paired end reads that failed previous minimum quality controls were discarded. Individual read groups were aligned, using bwa v0.7.12 with default parameters, to the UCSC hg19 reference human genome from Illumina iGenomes web site. Trimming rates and insert length were controlled on each read group based on metrics reported by Trimmomatic, and Picard v1.128 respectively.

Aligned reads from multiple read groups belonging to the same sample were indexed, sorted and merged using sambamba v0.5.4. Amplification duplicates were removed using Picard.

Various quality controls parameters were used including the obtained target coverage of the Nextera Rapid Capture exome library v1.2, mapping rates and duplication rates, based on metrics collected for each sample using Samtools v1.2, Picard v1.141, bedtools v2.25.0, and aggregated using custom Python v2.7.9 codes.

We applied GATK v3.5.0 base quality score recalibration and indel realignment [14] with standard parameters. We performed SNP and INDEL discovery and genotyping across each cohort of samples simultaneously using standard hard filtering parameters according to GATK Best Practices recommendations.

All variants were annotated with functional prediction using SnpEff v4.2. Additionally, functional annotation of variants found in two public databases (NCBI dbSNP v144 and dbNSFP v2.9) was added using SnpSift, part of the same software package. Multiallelic variants were decomposed and normalized using vt. A GEMINI v0.18.3; a database was created [15], and variants selected according to functional rules. Finally, they were manually validated against read alignments, using Integrative Genomics Viewer software (IGV) v2.3. Coverage metrics were calculated using bedtools.

## Supplementary Data

S1. Details of study samples. Table 1 Clinical data of the PPOLs and HNSCC cell lines/cultures; Table 2 Matching fibroblasts for PPOLs and HNSCC cell lines/cultures; Table 3 Primary HNSCC and matching normal mucosa samples used for pyrosequencing methylation analyses and gene expression analyses.

S2A. Cumulative distribution of coverage showing fraction of targeted bsed (Y-axis) that’s were covered by at least certain depth (x-axis) in exome sequencing

S2B. Cumulative distribution of coverage showing fraction of targeted bsed (Y-axis) that’s were covered by at least certain depth (x-axis) in HaloPlex sequencing

S2C. Significant variants in HNSCC driver genes from exome sequencing.

S2D. Significant variants in HNSCC driver genes from HaloPlex sequencing.

S3. Frequency of copy number alterations in IntOgen-derived cancer driver genes PPOLs and HNSCC cell lines. Table 1: Cancer drivers showing low copy number gains (≤ 2) ordered by frequency in LN+ve HNSCC cell lines with minimum frequency of 40%; Table 2: Cancer drivers showing low copy number gains (≤ 2) ordered by frequency in LN-ve HNSCC cell lines with minimum frequency of 40%; Table 3: Cancer drivers showing high copy number gains (>2) ordered by frequency in LN+ve HNSCC cell lines with minimum frequency of 10%; Table 4: Cancer drivers showing high copy number gains (>2) ordered by frequency in LN-ve HNSCC cell lines with minimum frequency of 10%; Table 5: Cancer drivers showing hemizygous loss ordered by frequency in LN+ve HNSCC cell lines with minimum frequency of 40%; Table 6: Cancer drivers showing hemizygous loss ordered by frequency in LN-ve HNSCC cell lines with minimum frequency of 40%; Table 7: Cancer drivers showing homozygous loss ordered by frequency in LN+ve HNSCC cell lines.

S4. Subtraction karyogram of mortal PPOLs and matched normal fibroblasts.

S5. Immortal PPOL cell lines SCNA density plots for chromosome 3, 8, 9 and 20.

S6. GISTIC peak regions in PPOL and HNSCC cell lines. Table 1. GISTIC peak regions in PPOLs with varying significance thresholds (Q bound value); Table 2. GISTIC peak regions in HNSCCs with varying significance thresholds (Q bound value); Table 3. GISTIC peak regions in LN-ve HNSCCs with Q bound value<0.25; Table 4. GISTIC peak regions in LN+ve HNSCCs with Q bound value<0.25.

S7. Analysis of *NCKAP5*. Table 1. Analysis of *NCKAP5* mutations listed in COSMIC Fig 1. Hemizygous and homozygous deletions at *NCKAP5* locus; Fig.2. Expression analyses of *NCKAP5* in HNSCC; Figure 3. Expression analyses of *NCKAP5* in other tumour types.

S8. Deletions in HNSCC cell lines at the *CSMD1* locus.

S9. Modulation of the *CSMD1* expression in HNSCC cell lines.

S10. Comparison of SCNA in two HNSCC panels.

S11. Integrative analyses of somatic copy number changes and gene expression. Table 1. Copy number gains in PPOLs and HNSCC of genes that show correlation with expression in SCNA regions showing significant difference between PPOLs and HNSCCs; Table 2. Copy number losses in PPOLs and HNSCC of genes that show correlation with expression in SCNA regions showing significant difference between PPOLs and HNSCCs.

S12. Hierarchical cluster analyses of immortal HNSCC cell lines for high copy number SCNA – association with lymph node metastases.

S13. KEGG pathway genes enrichment in GISTIC peak regions

S14. Analysis of CLDN1 and BCL2L1 in primary HNSCC and PPOL. Fig 1. Expression of CLDN1 in PPOLS and HNSC; Figure 2. Expression of BCL-XL in PPOLS and HNSCC

S15. SCNAs of genes in cancer-related KEGG pathway enriched in GISTIC regions. Fig 1 Pathway in cancer; Fig 2 Apoptosis pathway; Fig 3 Axon guidance pathway; Fig 4 Cell adhesion pathway; Fig 5 Cell cycle pathway; Fig 6 Endocytosis pathway; Fig 7. JAK-Stat pathway; Fig 8 MAPK pathway; Fig 9 Ubuiquitin-proteosome pathway; Fig 10 WNT pathway.

S16A. Somatic copy number changes in *ADAMTS9* in a multitumour cell line panel.

S16B. Analyses of *ADAMTS9* sequence variants identified in this study reported in Stransky et al., 2011 and COSMIC database.

S17. Gene size and density for genes sustaining homozygous deletions in immortal HNSCC cell lines.

## References

1. Ferlay J, Soerjomataram I, Dikshit R, Eser S, Mathers C, Rebelo M, et al. Cancer incidence and mortality worldwide: sources, methods and major patterns in GLOBOCAN 2012. Int J Cancer. 2015;136(5):E359–86.

2. Pace-Balzan A, Shaw RJ, Butterworth C. Oral rehabilitation following treatment for oral cancer. Periodontol 2000. 2011;57(1):102–17.

3. Stransky N, Egloff AM, Tward AD, Kostic AD, Cibulskis K, Sivachenko A, et al. The mutational landscape of head and neck squamous cell carcinoma. Science. 2011;333(6046):1157–60.

4. Agrawal N, Frederick MJ, Pickering CR, Bettegowda C, Chang K, Li RJ, et al. Exome sequencing of head and neck squamous cell carcinoma reveals inactivating mutations in NOTCH1. Science. 2011;333(6046):1154–7.

5. Pickering CR, Zhang J, Yoo SY, Bengtsson L, Moorthy S, Neskey DM, et al. Integrative genomic characterization of oral squamous cell carcinoma identifies frequent somatic drivers. Cancer Discov. 2013;3(7):770–81.

6. India Project Team of the International Cancer Genome C. Mutational landscape of gingivo-buccal oral squamous cell carcinoma reveals new recurrently-mutated genes and molecular subgroups. Nat Commun. 2013;4:2873.

7. The Cancer Genome Atlas N. Comprehensive genomic characterization of head and neck squamous cell carcinomas. Nature. 2015;517(7536):576–82.

8. Bhattacharya A, Roy R, Snijders AM, Hamilton G, Paquette J, Tokuyasu T, et al. Two distinct routes to oral cancer differing in genome instability and risk for cervical node metastasis. Clin Cancer Res. 2011;17(22):7024–34.

9. Hunter KD, Parkinson EK, Harrison PR. Profiling early head and neck cancer. Nat Rev Cancer. 2005;5(2):127–35.

10. Ha PK, Benoit NE, Yochem R, Sciubba J, Zahurak M, Sidransky D, et al. A transcriptional progression model for head and neck cancer. Clin Cancer Res. 2003;9(8):3058–64.

11. Hunter KD, Thurlow JK, Fleming J, Drake PJ, Vass JK, Kalna G, et al. Divergent routes to oral cancer. Cancer Res. 2006;66(15):7405–13.

12. Bedi GC, Westra WH, Gabrielson E, Koch W, Sidransky D. Multiple head and neck tumors: evidence for a common clonal origin. Cancer Res. 1996;56(11):2484–7.

13. Tabor MP, Brakenhoff RH, Ruijter-Schippers HJ, Kummer JA, Leemans CR, Braakhuis BJ. Genetically altered fields as origin of locally recurrent head and neck cancer: a retrospective study. Clin Cancer Res. 2004;10(11):3607–13.

14. Califano J, van der Riet P, Westra W, Nawroz H, Clayman G, Piantadosi S, et al. Genetic progression model for head and neck cancer: implications for field cancerization. Cancer Res. 1996;56(11):2488–92.

15. Edington KG, Loughran OP, Berry IJ, Parkinson EK. Cellular immortality: a late event in the progression of human squamous cell carcinoma of the head and neck associated with p53 alteration and a high frequency of allele loss. Mol Carcinog. 1995;13(4):254–65.

16. McGregor F, Muntoni A, Fleming J, Brown J, Felix DH, MacDonald DG, et al. Molecular changes associated with oral dysplasia progression and acquisition of immortality: potential for its reversal by 5-azacytidine. Cancer Res. 2002;62(16):4757–66.

17. Rheinwald JG, Beckett MA. Tumorigenic keratinocyte lines requiring anchorage and fibroblast support cultured from human squamous cell carcinomas. Cancer Res. 1981;41(5):1657–63.

18. Wood HM, Daly C, Chalkley R, Senguven B, Ross L, Egan P, et al. The genomic road to invasion-examining the similarities and differences in the genomes of associated oral pre-cancer and cancer samples. Genome Med. 2017;9(1):53.

19. Muntoni A, Fleming J, Gordon KE, Hunter K, McGregor F, Parkinson EK, et al. Senescing oral dysplasias are not immortalized by ectopic expression of hTERT alone without other molecular changes, such as loss of INK4A and/or retinoic acid receptor-beta: but p53 mutations are not necessarily required. Oncogene. 2003;22(49):7804–8.

20. McGregor F, Wagner E, Felix D, Soutar D, Parkinson K, Harrison PR. Inappropriate Retinoic Acid Receptor-β Expression in Oral Dysplasias: Correlation with Acquisition of the Immortal Phenotype. Cancer Research. 1997;57(18):3886–9.

21. Paila U, Chapman BA, Kirchner R, Quinlan AR. GEMINI: Integrative Exploration of Genetic Variation and Genome Annotations. PLoS Comput Biol. 2013;9(7):e1003153.

22. Lawrence MS, Stojanov P, Polak P, Kryukov GV, Cibulskis K, Sivachenko A, et al. Mutational heterogeneity in cancer and the search for new cancer-associated genes. Nature. 2013;499(7457):214–8.

23. El-Naggar AK, Lai S, Luna MA, Zhou X-D, Weber RS, Goepfert H, et al. Sequential p53 mutation analysis of pre-invasive and invasive head and neck squamous carcinoma. International Journal of Cancer. 1995;64(3):196–201.

24. Papadimitrakopoulou Vali IJ, Lippman M Scott, Lee Soo Jin, Fan Hong You, Clayman Gary, Ro Y Jay, Hittelman N Walter, Lotan Reuben, Hong K Waun and Mao Li Frequent inactivation of p16INK4a in oral premalignant lesions. Oncogene. 1997;14:1799–803.

25. Qin G-Z, Park JY, Chen S-Y, Lazarus P. A high prevalence of p53 mutations in pre-malignant oral erythroplakia. International Journal of Cancer. 1999;80(3):345–8.

26. Beroukhim R, Getz G, Nghiemphu L, Barretina J, Hsueh T, Linhart D, et al. Assessing the significance of chromosomal aberrations in cancer: Methodology and application to glioma. Proceedings of the National Academy of Sciences. 2007;104(50):20007–12.

27. Kluth M, Galal R, Krohn A, Weischenfeldt J, Tsourlakis C, Paustian L, et al. Prevalence of chromosomal rearrangements involving non-ETS genes in prostate cancer. International journal of oncology. 2015;46(4):1637–42.

28. Buday L, Wunderlich L, Tamas P. The Nck family of adapter proteins: regulators of actin cytoskeleton. Cell Signal. 2002;14(9):723–31.

29. Joannes A, Bonnomet A, Bindels S, Polette M, Gilles C, Burlet H, et al. Fhit regulates invasion of lung tumor cells. Oncogene. 2010;29(8):1203–13.

30. Roz L, Gramegna M, Ishii H, Croce CM, Sozzi G. Restoration of fragile histidine triad (FHIT) expression induces apoptosis and suppresses tumorigenicity in lung and cervical cancer cell lines. Proc Natl Acad Sci U S A. 2002;99(6):3615–20.

31. Beroukhim R, Mermel CH, Porter D, Wei G, Raychaudhuri S, Donovan J, et al. The landscape of somatic copy-number alteration across human cancers. Nature. 2010;463(7283):899–905.

32. Shull AY, Clendenning ML, Ghoshal-Gupta S, Farrell CL, Vangapandu HV, Dudas L, et al. Somatic mutations, allele loss, and DNA methylation of the Cub and Sushi Multiple Domains 1 (CSMD1) gene reveals association with early age of diagnosis in colorectal cancer patients. PloS one. 2013;8(3):e58731.

33. Wilhelm M, Schlegl J, Hahne H, Gholami AM, Lieberenz M, Savitski MM, et al. Mass-spectrometry-based draft of the human proteome. Nature. 2014;509(7502):582–7.

34. Gross AM, Orosco RK, Shen JP, Egloff AM, Carter H, Hofree M, et al. Multi-tiered genomic analysis of head and neck cancer ties TP53 mutation to 3p loss. Nat Genet. 2014;46(9):939–43.

35. Lo PH, Leung AC, Kwok CY, Cheung WS, Ko JM, Yang LC, et al. Identification of a tumor suppressive critical region mapping to 3p14.2 in esophageal squamous cell carcinoma and studies of a candidate tumor suppressor gene, ADAMTS9. Oncogene. 2007;26(1):148–57.

36. Lung HL, Lo PH, Xie D, Apte SS, Cheung AK, Cheng Y, et al. Characterization of a novel epigenetically-silenced, growth-suppressive gene, ADAMTS9, and its association with lymph node metastases in nasopharyngeal carcinoma. Int J Cancer. 2008;123(2):401–8.

37. Lo PH, Lung HL, Cheung AK, Apte SS, Chan KW, Kwong FM, et al. Extracellular protease ADAMTS9 suppresses esophageal and nasopharyngeal carcinoma tumor formation by inhibiting angiogenesis. Cancer Res. 2010;70(13):5567–76.

38. Forbes SA, Beare D, Gunasekaran P, Leung K, Bindal N, Boutselakis H, et al. COSMIC: exploring the world’s knowledge of somatic mutations in human cancer. Nucleic Acids Research. 2015;43(D1):D805–D11.

39. Qi J, Yu Y, Akilli Öztürk Ö, Holland JD, Besser D, Fritzmann J, et al. New Wnt/β-catenin target genes promote experimental metastasis and migration of colorectal cancer cells through different signals. Gut. 2015.

40. Chung K-Y, Cheng IKC, Ching AKK, Chu J-H, Lai PBS, Wong N. Block of proliferation 1 (BOP1) plays an oncogenic role in hepatocellular carcinoma by promoting epithelial-to-mesenchymal transition. Hepatology. 2011;54(1):307–18.

41. Reed AL, Califano J, Cairns P, Westra WH, Jones RM, Koch W, et al. High Frequency of p16 (CDKN2/MTS-1/INK4A) Inactivation in Head and Neck Squamous Cell Carcinoma. Cancer Research. 1996;56(16):3630–3.

42. Cairns P, Polascik TJ, Eby Y, Tokino K, Califano J, Merlo A, et al. Frequency of homozygous deletion at p16/CDKN2 in primary human tumours. Nat Genet. 1995;11(2):210–2.

43. Munro J, Stott FJ, Vousden KH, Peters G, Parkinson EK. Role of the alternative INK4A proteins in human keratinocyte senescence: evidence for the specific inactivation of p16INK4A upon immortalization. Cancer Res. 1999;59(11):2516–21.

44. Castets M, Broutier L, Molin Y, Brevet M, Chazot G, Gadot N, et al. DCC constrains tumour progression via its dependence receptor activity. Nature. 2012;482(7386):534–7.

45. Escudero-Esparza A, Bartoschek M, Gialeli C, Okroj M, Owen S, Jirstrom K, et al. Complement inhibitor CSMD1 acts as tumor suppressor in human breast cancer. Oncotarget. 2016;7(47):76920–33.

46. Tang MR, Wang YX, Guo S, Han SY, Wang D. CSMD1 exhibits antitumor activity in A375 melanoma cells through activation of the Smad pathway. Apoptosis. 2012;17(9):927–37.

47. Liesenfeld M, Mosig S, Funke H, Jansen L, Runnebaum IB, Dürst M, et al. SORBS2 and TLR3 induce premature senescence in primary human fibroblasts and keratinocytes. BMC Cancer. 2013;13(1):1–11.

48. Patel K, Scrimieri F, Ghosh S, Zhong J, Kim M-S, Ren YR, et al. FAM190A Deficiency Creates a Cell Division Defect. The American Journal of Pathology. 2013;183(1):296–303.

49. Bashaw GJ, Klein R. Signaling from Axon Guidance Receptors. Cold Spring Harbor Perspectives in Biology. 2010;2(5).

50. Huang M, Anand S, Murphy EA, Desgrosellier JS, Stupack DG, Shattil SJ, et al. EGFR-dependent pancreatic carcinoma cell metastasis through Rap1 activation. Oncogene. 2012;31(22):2783–93.

51. Errington TM, Macara IG. Depletion of the adaptor protein NCK increases UV-induced p53 phosphorylation and promotes apoptosis. PloS one. 2013;8(9):e76204.

52. Kresty LA, Mallery SR, Knobloch TJ, Song H, Lloyd M, Casto BC, et al. Alterations of p16(INK4a) and p14(ARF) in patients with severe oral epithelial dysplasia. Cancer Res. 2002;62(18):5295–300.

53. Malkoski SP, Wang X-J. Two sides of the story? Smad4 loss in pancreatic cancer versus head-and-neck cancer. FEBS Letters. 2012;586(14):1984–92.

54. Huntley SP, Davies M, Matthews JB, Thomas G, Marshall J, Robinson CM, et al. Attenuated type II TGF-β receptor signalling in human malignant oral keratinocytes induces a less differentiated and more aggressive phenotype that is associated with metastatic dissemination. International Journal of Cancer. 2004;110(2):170–6.

55. Alani RM, Hasskarl J, Grace M, Hernandez M-C, Israel MA, Münger K. Immortalization of primary human keratinocytes by the helix–loop–helix protein, Id-1. Proceedings of the National Academy of Sciences. 1999;96(17):9637–41.

56. Lin J, Guan Z, Wang C, Feng L, Zheng Y, Caicedo E, et al. Inhibitor of Differentiation 1 Contributes to Head and Neck Squamous Cell Carcinoma Survival via the NF-κB/Survivin and Phosphoinositide 3-Kinase/Akt Signaling Pathways. American Association for Cancer Research. 2010;16(1):77–87.

57. Burns JE, Clark LJ, Yeudall WA, Mitchell R, Mackenzie K, Chang SE, et al. The p53 status of cultured human premalignant oral keratinocytes. Br J Cancer. 1994;70(4):591–5.

58. Sun Q, Wang R, Luo J, Wang P, Xiong S, Liu M, et al. Notch1 promotes hepatitis B virus X protein-induced hepatocarcinogenesis via Wnt/beta-catenin pathway. Int J Oncol. 2014;45(4):1638–48.

59. Lin JT, Chen MK, Yeh KT, Chang CS, Chang TH, Lin CY, et al. Association of high levels of Jagged-1 and Notch-1 expression with poor prognosis in head and neck cancer. Ann Surg Oncol. 2010;17(11):2976–83.

60. Song X, Xia R, Li J, Long Z, Ren H, Chen W, et al. Common and complex Notch1 mutations in Chinese oral squamous cell carcinoma. Clin Cancer Res. 2014;20(3):701–10.

61. Lefort K, Mandinova A, Ostano P, Kolev V, Calpini V, Kolfschoten I, et al. Notch1 is a p53 target gene involved in human keratinocyte tumor suppression through negative regulation of ROCK1/2 and MRCKalpha kinases. Genes Dev. 2007;21(5):562–77.

62. Sakamoto K, Fujii T, Kawachi H, Miki Y, Omura K, Morita K, et al. Reduction of NOTCH1 expression pertains to maturation abnormalities of keratinocytes in squamous neoplasms. Lab Invest. 2012;92(5):688–702.

63. Nickoloff BJ, Qin JZ, Chaturvedi V, Denning MF, Bonish B, Miele L. Jagged-1 mediated activation of notch signaling induces complete maturation of human keratinocytes through NF-kappaB and PPARgamma. Cell Death Differ. 2002;9(8):842–55.

64. Lowell S, Jones P, Le Roux I, Dunne J, Watt FM. Stimulation of human epidermal differentiation by delta-notch signalling at the boundaries of stem-cell clusters. Curr Biol. 2000;10(9):491–500.

65. Demehri S, Turkoz A, Kopan R. Epidermal Notch1 loss promotes skin tumorigenesis by impacting the stromal microenvironment. Cancer Cell. 2009;16(1):55–66.

66. Patel V, Rosenfeldt HM, Lyons R, Servitja J-M, Bustelo XR, Siroff M, et al. Persistent activation of Rac1 in squamous carcinomas of the head and neck: evidence for an EGFR/Vav2 signaling axis involved in cell invasion. Carcinogenesis. 2007;28(6):1145–52.

67. Cerami E, Gao J, Dogrusoz U, Gross BE, Sumer SO, Aksoy BA, et al. The cBio Cancer Genomics Portal: An Open Platform for Exploring Multidimensional Cancer Genomics Data. Cancer Discovery. 2012;2(5):401–4.

68. Silvera D, Arju R, Darvishian F, Levine PH, Zolfaghari L, Goldberg J, et al. Essential role for eIF4GI overexpression in the pathogenesis of inflammatory breast cancer. Nat Cell Biol. 2009;11(7):903–8.

69. Lin C-J, Cencic R, Mills JR, Robert F, Pelletier J. c-Myc and eIF4F Are Components of a Feedforward Loop that Links Transcription and Translation. Cancer Research. 2008;68(13):5326–34.

70. Krauthammer M, Kong Y, Ha BH, Evans P, Bacchiocchi A, McCusker JP, et al. Exome sequencing identifies recurrent somatic RAC1 mutations in melanoma. Nat Genet. 2012;44(9):1006–14.

71. Wallingford JB, Habas R. The developmental biology of Dishevelled: an enigmatic protein governing cell fate and cell polarity. Development. 2005;132(20):4421–36.

72. Rudy SF, Brenner JC, Harris JL, Liu J, Che J, Scott MV, et al. In vivo Wnt pathway inhibition of human squamous cell carcinoma growth and metastasis in the chick chorioallantoic model. Journal of Otolaryngology-Head & Neck Surgery. 2016;45(1):1–8.

73. Yau C, Mouradov D, Jorissen RN, Colella S, Mirza G, Steers G, et al. A statistical approach for detecting genomic aberrations in heterogeneous tumor samples from single nucleotide polymorphism genotyping data. Genome Biol. 2010;11(9):R92.

